# Design of the mammalian cone photoreceptor to Off bipolar cell synapse

**DOI:** 10.1101/2022.03.04.483056

**Authors:** Chad P. Grabner, Daiki Futagi, Jun Shi, Vytas Bindokas, Katsunori Kitano, Eric Schwartz, Steven H. DeVries

## Abstract

Graded synapses in sensory systems reliably transmit small signals in the presence of continuous quantal noise. To understand how signaling is optimized during graded transmission, we counted the number of vesicles released by a mammalian cone terminal and compared it to the simultaneous responses in each Off bipolar cell type. Off bipolar cells contacting the terminal base comprised two groups depending on how they sampled transmitter release. In both groups, responses initially grew non-linearly with the number of released vesicles implicating a role for cooperativity during sparse release. One group sampled release from most of a cone’s ∼20 ribbons and can exploit averaging to improve signal reliability. The other, less-sensitive group made 1-3 contacts at the terminal center and responded to pooled transmitter, a consequence of membrane depolarization, using an insensitive kainate receptor. Off bipolar cells use different strategies to minimize transmission noise and encode cone output over different ranges.

## Introduction

The cone photoreceptor synapse is an essential link in the pathway that mediates daylight vision. At the synapse, graded changes in membrane voltage are encoded at ∼20 ribbon release sites (Chun et al., 1996; Herr et al., 2003; Zhang et al., 2020) by a spatiotemporal pattern of vesicle fusion. The resulting glutamate concentration profile in the cone synaptic cleft is then sampled and re-encoded as electrical signals by the dendritic contacts of more than a dozen types of postsynaptic On and Off bipolar cells (Wassle et al., 2009; Light et al., 2012; Helmstaedter et al., 2013; Tsukamoto and Omi, 2015; Shekhar et al., 2016; Tsukamoto and Omi, 2016; Franke et al., 2017). Transmitter quantization and resampling can introduce noise and potentially limit visual performance at graded synapses. However, in dim to moderate light, where the mean rates of vesicle release are relatively high (Choi et al., 2005), visual responses in the On pathways are limited by noise in cone phototransduction (Angueyra and Rieke, 2013) rather than by noise at the cone synapse or from other retinal sources (Ala-Laurila et al., 2011). Noise due to transmitter quanta in the On bipolar cell pathway may be suppressed by a metabotropic receptor-driven cascade (Masu et al., 1995) that can act as a low-pass filter (Ashmore and Copenhagen, 1980). Whether quantal noise affects transmission to Off bipolar cells, which, respond to glutamate release with AMPA/kainate (DeVries, 2000) receptors, is less clear. In support of a limiting effect of quantal noise on transmission in Off pathways, measurements in horizontal cells, which are postsynaptic to cones and express AMPA receptors, show an unexpected decrement in sensitivity in moderate light (Borghuis et al., 2009). In addition, quantal noise may have a greater impact on signal transmission in bright backgrounds where cones are hyperpolarized (Borghuis et al., 2018).

Membrane hyperpolarization slows the rate of vesicle fusion (Choi *et al*., 2005), providing a sparse substrate for signaling under the same conditions where transduction noise is reduced and flash responses are attenuated and shortened by adaptation (Angueyra and Rieke, 2013; Borghuis *et al*., 2018). Little is known about the mechanisms that optimize transmission between cones and postsynaptic Off bipolar cells, especially during strong background illumination.

Mammalian photoreceptor terminals contain two specialized synaptic structures: invaginations, which are found in both rods and cones, and basal junctions, which are unique to cones (Dowling and Boycott, 1966; Dowling, 2012). Each cone terminal contains approximately twenty, 200-400 nm deep membrane invaginations that open onto a compact terminal base (Missotten, 1965). Invaginations are marked at their apices by an intracellular ribbon surrounded by synaptic vesicles. Where ribbons approach the plasmamembrane, a close association between sustained, L-type Ca^2+^ channels and docked vesicles enables a stream of fusion events at a rate that is related to both membrane voltage (Jarsky et al., 2010; Bartoletti et al., 2011) and the complex kinetics of vesicle replenishment (Jackman et al., 2009; Mehta et al., 2013). In mammals, with few exceptions (West, 1976; Behrens et al., 2016), the dendrites of On bipolar cells enter invaginations and terminate centrally in relation to release sites (Kolb, 1970; Famiglietti and Kolb, 1976). At the terminal base, cone and bipolar cell membranes abut across a narrow, 13-16 nm (Lasansky, 1969; Raviola and Gilula, 1975) cleft to form basal junctions. At basal junctions, the cone membrane lacks the specializations that are typically associated with transmitter release including clusters of vesicles, active zones, and vesicle fusion profiles (Raviola and Gilula, 1975; Schaeffer et al., 1982). There is also no evidence for ectopic release (Van Hook and Thoreson, 2015). Rather, to reach basal contacts, glutamate diffuses from ribbon sites over a 200 – 750 nm extracellular path. In mammals, Off bipolar cells almost exclusively contact cones at basal junctions (Hopkins and Boycott, 1997). An obligatory diffusion distance from ribbon release sites to basal junctions may challenge the classical notion that individual quanta are the basis for synaptic transmission. Additionally, the location of basal contacts between invaginations raises the possibility that a dendritic contact can be excited by near simultaneous release events from several nearby invaginations.

At least two mechanisms have been proposed to improve the ratio of signal to noise (SNR) at photoreceptor synapses. In the first mechanism, established at photoreceptor to second order neuron synapses in Drosophila, the SNR is improved by sampling from most or all output ribbons. Since each ribbon signals photoreceptor membrane voltage independently and release at a ribbon is stochastic, the signal increases in proportion to the number of ribbons sampled, *n*, while the noise of transmission, expressed as a standard deviation, increases only as √*n* (de Ruyter van Steveninck and Laughlin, 1996; Laughlin et al., 1998). Ultrastructural studies on cone to midget bipolar synapses, components of specialized pathways in the primate retina, show a dense innervation that is well-suited to sample release from all of a cone’s ribbons (Tsukamoto and Omi, 2015; 2016; Zhang *et al*., 2020), but a functional demonstration in mammals is lacking. The second mechanism, described at the mammalian rod to On bipolar cell synapse, involves a threshold non-linearity that is optimally placed to filter out a continuous low-amplitude noise that originates in phototransduction while transmitting larger amplitude signals that result from most photoresponses (Field and Rieke, 2002). The threshold is produced by a saturation in the On bipolar cell transduction cascade, which occurs when the concentration of cleft glutamate is high during continuous vesicle fusion in the dark (Sampath and Rieke, 2004). Conceptually, for Off bipolar cells, a “threshold” non-linearity might reduce the postsynaptic effect of individual transmitter quanta, whose arrival times under steady conditions are random, while favoring a response to multivesicular release, which can be coordinated by a change in membrane voltage. A recent report demonstrates a non-linearity at the synapses between cones and some Off bipolar cells in the salamander retina (Schreyer and Gollisch, 2021).

To address the mechanisms of signaling at the cone to Off bipolar cell basal synapse, we briefly depolarized a ground squirrel cone in voltage clamp and counted the number of released vesicles by measuring the current response of the cone terminal glutamate transporter (Szmajda and Devries, 2011). At the same time, we recorded the epsc response in identified postsynaptic Off bipolar cells (DeVries et al., 2006). Additionally, we took advantage of a design feature of the ground squirrel cone synapse that is unique among mammalian retinas: The Off cb2 bipolar cell makes invaginating or synaptic ribbon-associated contacts (Grabner et al., 2016). This feature enabled us to compare the effect of synaptic location, invaginating versus basal, on signaling within the Off bipolar cell types. We found that cb2 cells, which contacted a subset of a cone’s invaginations, sampled a constant fraction of a cone’s released vesicles starting from the uniquantal level. In contrast, basally contacting cb3 and cb1 cells displayed threshold nonlinearities, detecting small release events minimally or not at all, respectively. Above threshold, cb3 cells, which make multiple cone contacts, used a sensitive kainate receptor to sample release from nearly all of a cone’s invaginations, and thus may employ a classical averaging strategy. Different from cb3 cells, cb1 cells expressed an insensitive kainate receptor and made only a few contacts in the center of the cone terminal. The properties of cb1 cell synapses are well-suited to signal tidal release, a consequence of event summation, which can accompany large membrane depolarizations. Thus, Off bipolar cell types use both thresholding and sampling strategies to optimize signal transmission while encoding the cone output over different but overlapping ranges.

## Results

Synaptic ribbons distributed across the concaved surfaces of two cone terminals are labeled with an antibody directed against the active zone protein bassoon (Dick et al., 2003) (Figure 1A). In the ground squirrel, most cones are contacted by all five Off bipolar cell types (Figure 1B) (Light *et al*., 2012). We used a STED super-resolution microscope to obtain 3D images of the dendrites of individually labeled cb1a, cb2, and cb3b cells at representative cone terminals (Figure 1C). In preliminary observations, each type contacted the cone terminal in a distinctive pattern. The dendrites of cb2 cells, which, along with horizontal cells, express GluA4 receptors (Pan and Massey, 2007; Lindstrom et al., 2014), ended near clusters of GluA4 subunits that were juxtaposed with the synaptic ribbon protein Ribeye. This association is consistent with an invaginating locus. The dendrites of cb1a cells contacted cones in the center of the terminal relatively distant from clusters of GluA4 receptors and ribbons. The dendrites of cb3b cells made multiple contacts that were also distant from clusters and ribbons. These initial observations which suggest that Off bipolar have different contact patterns prompted us to determine whether the Off bipolar cell types sampled the population of ribbon release sites in different ways.

**Figure 1.**
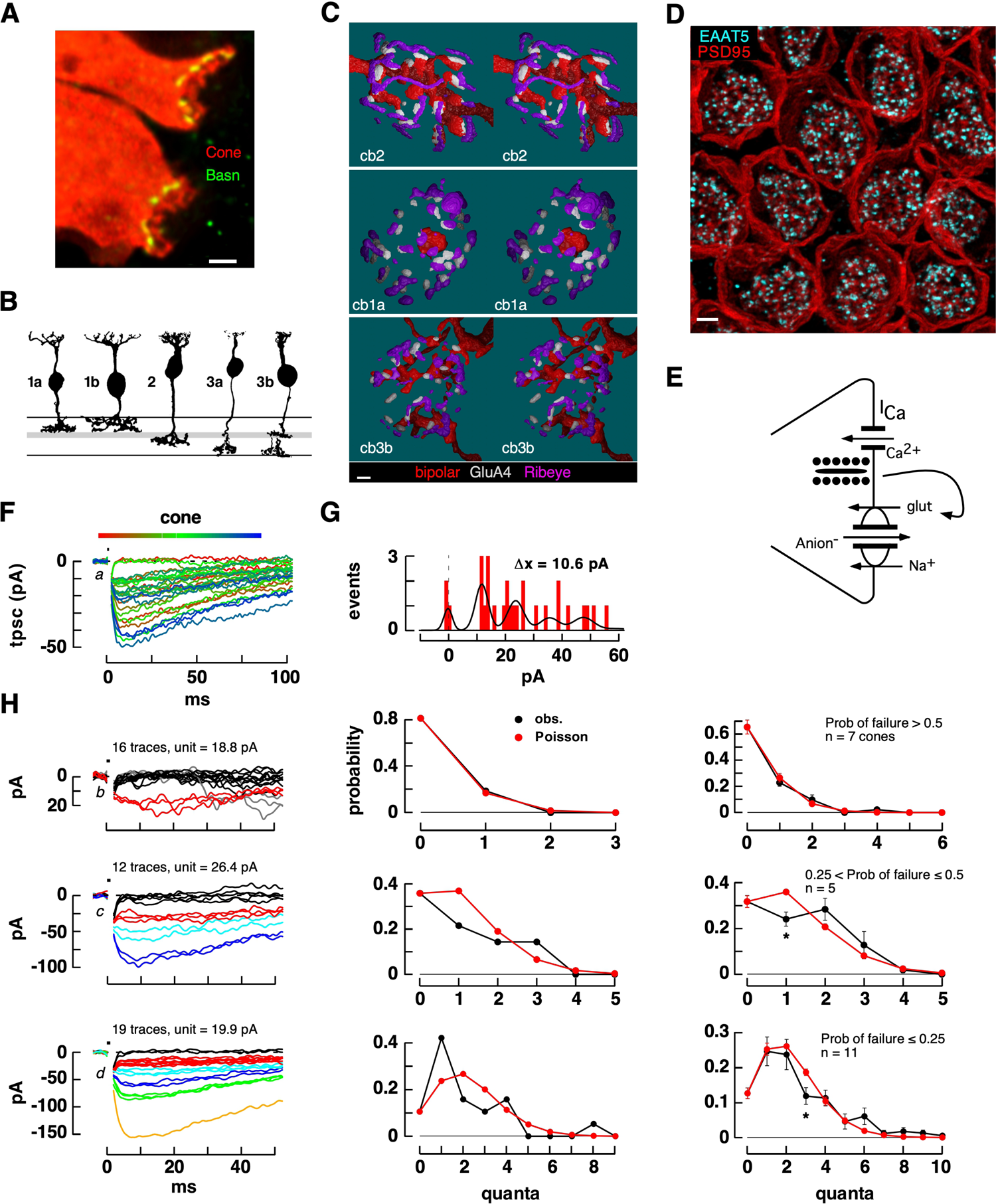
Cone transporters signal evoked glutamate release. **A.** Cross-sectional thin-stack STED image of neighboring cones labeled with antibodies to the tracer Cascade Blue (red) and bassoon (green). Scale = 1 µm. **B.** Diagram of the five main Off bipolar cell types in the ground squirrel. **C.** Stereo images of 3D reconstructions of individual cone terminals labeled with antibodies to Ribeye (magenta), GluA4 (white), and GFP (red) that was virally transduced into bipolar cells. Scale = 0.5 µm. **D.** Transporter clusters at the base of the cone terminal. Wholemount STED image of cone terminals immunolabeled for PSD95, a marker for the sides and base of the terminal (red), and EAAT5 (cyan), localized to the base (scale bar = 2 µm). **E.** Diagram showing the activation of a transporter anion conductance by released glutamate. **F.** A train of eighteen 1 ms pulses from −70 to −30 mV (black mark, above) elicited consecutive transporter current responses. Responses are color-coded according to temporal position in the sequence (horizontal bar, above). Failures and successes were interspersed. The long duration of the transporter response likely reflects the slow turnover rate of EAAT5 rather than a persistent elevation of cleft glutamate (Gameiro *et al*., 2011). **G**. Amplitude histogram of the responses in F fitted with a sum-of-Gaussians (interpeak distance = 10.56 pA). **H.** Comparison between observed and predicted amplitude distributions. Predicted distributions were calculated from the measured failure probability, P_0_, ranging from high (top row), to medium (middle), to low (bottom). Left column: consecutive tpsc responses during a train of steps to the same voltage color-coded according to amplitude histogram-defined quantal levels. Responses from 3 different cones. Center column: observed and predicted quantal content probabilities for the responses in the left column. Right column: aggregate data. Responses were grouped and averaged according to ranges of P_0_. Asterisks denote values that were significantly different at the p = 0.01 – 0.05 level. Letters *a-d* refer to Fig S1B.

### Cones report vesicle-released glutamate

Released glutamate feeds back onto the presynaptic cone terminal to elicit a long-lasting inward current that is mediated by an anion conductance intrinsic to a TBOA-sensitive glutamate transporter (Fig. 1E) (Szmajda and Devries, 2011). An antibody to the glutamate transporter EAAT5 strongly labels mouse cone terminals (Gehlen et al., 2021), and STED images of the ground squirrel terminal show distributed EAAT5 clusters at the base (Fig. 1D; supplemental movie 1; 62.0 ± 5.4 puncta per cone, mean ± S.D., n = 9 cones). We used the cone transporter presynaptic current (tpsc) to obtain an approximate count of the vesicles released by a cone by applying brief, depolarizing pulses from −70 to −30 mV at a rate of 1-2 Hz. In a typical experiment, a train of pulses elicited cone tpscs that were interspersed with failures (Fig 1F). An estimate of the unitary event amplitude, 10.6 pA, was obtained by fitting an amplitude histogram with a sum of Gaussians (Fig 1G; 15.5 ± 5.0 pA, mean ± S.D. in 24 cones, Fig S1A). Spontaneous events have a similar amplitude (Szmajda and Devries, 2011). Tpsc amplitudes were Poisson distributed (Fig 1H), especially during steps to relatively hyperpolarized voltages that were marked by a high probability of response failure, P_0_ (Fig 1H, top row). Deviations from the Poisson prediction were more evident during stronger depolarizations that had a lower percentage of response failures such that there was a slight deficit in small events and an excess in large events (Fig 1H, middle and bottom rows). The skew in observed responses towards larger events could be due to multiquantal release (Singer et al., 2004). To estimate and effective unitary event size, we relied on amplitude histograms and measurements of P_0_ rather than the response variance to mean ratio, which can be dramatically altered by the presence of multiquantal events (Fig S1B). Plots of the variance to mean ratio as a function of mean response amplitude were nonetheless useful for demonstrating transporter saturation, which began when tpscs exceeded approximately 35% of their maximal value (Fig S1C) or 150 – 200 pA (roughly 10 – 15 released units from an estimated 20 ribbons). An onset for transporter saturation at low levels of release is unsurprising given an EC_50_ of ∼60 µM for EAAT5 (Gameiro et al., 2011) and the distribution of transporters at the cone base.

### Sampling of glutamate release by invaginating cb2 Off bipolar cells

We used the cone to cb2 cell synapse to test our paired recording approach. On the idea that receptors on invaginating cb2 cell dendritic tips exclusively respond to vesicle release at the local ribbon, we hypothesized a linear relationship between cone tpsc and bipolar cell epsc amplitudes with a slope determined by the fraction of release sites sampled and an intercept near the origin. Stepping a cone from −70 to −20 mV for 1 ms produced maximal peak responses of −457 and − 659 pA in the cone and bipolar cell, respectively (Fig 2A). We then applied a sequence of 1 ms pulses from −70 to −30 mV to the cone and recorded the current responses in the cone and bipolar cell. Responses were grouped into three sets for illustration. First, response failures in the cone were always associated with failures in the cb2 bipolar cell (Fig 2B, top row). The opposite behavior, namely, bipolar cell events during cone response failures, would suggest incomplete reporting by the cone terminal. Second, we selected for the simultaneous occurrence of a cone response and a bipolar cell failure (Fig 2B, middle row). Cone responses and bipolar cell failures can occur if some cone release events are not detected by the postsynaptic cb2 cell. Evidence for postsynaptic sensitivity to individual fusion events comes from traces obtained under conditions of evoked release (Fig 2C) in which the simultaneous pre- and postsynaptic responses consisted of small events with stereotyped amplitudes. Cb2 cell responses to single packets of cone transmitter suggest that the absence of a bipolar cell response follows from an absence of local release rather than an inability to respond to single events. Finally, larger cone presynaptic responses were invariably associated with postsynaptic responses (Fig 2B, lower row). The occasional failure to detect release is consistent with the idea that a cb2 cell does not contact all of a cone’s invaginations and responds only to local quantal release.

**Figure 2.**
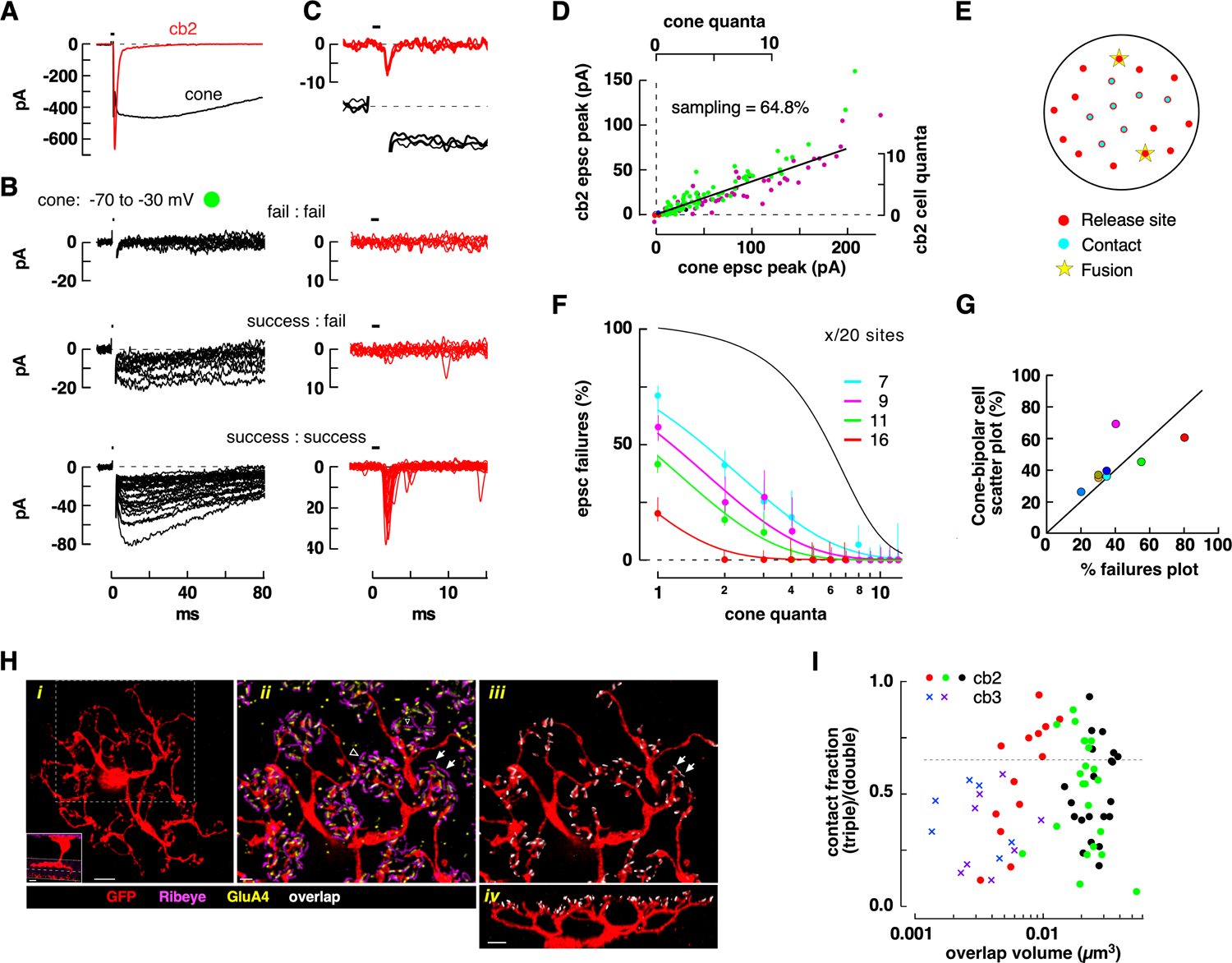
Linear responses of cb2 cells at the cone synapse. **A.** Maximal pre- and postsynaptic currents elicited by a brief cone membrane depolarization (horizontal bar). The bipolar cell was maintained at −70 mV. **B.** Grouped traces during a train of 78 consecutive cone steps from −70 to −30 mV. <u>Upper row</u>, *left*. Superimposed response failures in the cone. *right*. Superimposed corresponding traces from the bipolar cell. <u>Middle row</u>. The criterion for selecting traces was a combination of a cone response and bipolar cell failure to respond. <u>Bottom row</u>. Larger cone tpscs accompanied by bipolar cell epscs. All the responses from the train that met the stated criteria are shown in the upper and middle rows. Half of the responses in the lower row are omitted for clarity. Green circle indicates the corresponding data points in D. **C.** Three selected trace pairs superimposed showing apparent unitary responses in the cone and bipolar cell. Time bases in B apply to A and C. **D.** Plot of peak bipolar cell epsc versus cone tpsc response amplitude (left and lower axes, plot omits data resulting from the largest cone responses including those in A; purple circles plot results from a subsequent train of steps to −35 mV). Axes (top and right) were scaled by the calculated unitary event amplitudes in the cone and bipolar cell. **E.** Cone terminal model for cb2 cell responses. **F.** Plot of percent epsc response failures versus tpsc quantal content on a log scale for 4 cone to cb2 cell pairs. Theoretical curves were obtained from calculations that assumed that x = 7, 9, 11, or 16 cb2 contacts out of a maximum of 20 sites and that vesicle release occurs independently at each site. A success occurred when a vesicle fused at a release site occupied by a cb2 dendrite and a failure otherwise. The black curve shows a limiting case in which a cb2 cell samples from all 20 sites and a success requires the release of two or more vesicles at any site; cooperative sampling from fewer sites moves the black curve to the right. Error bars are statistical (see Methods). **H.** *(i).* (*inset*) Reconstructed cross-sectional confocal image of a GFP labeled cb2 cell with the axon arborizing in the top half of the IPL (scale = 5 µm). Maximum intensity projection from a 3D STED image stack of the labeled cb2 cell (scale = 3 µm). The region within the dashed boundary is shown in *ii* and *iii*. *(ii)*. Dendritic tree (red) with labelled Ribeye (magenta) and GluA4 (yellow). An open arrowhead shows a close association between Ribeye and GluA4 labeling without a nearby cb2 dendritic process (*i.e.*, a ‘double’) whereas the closed arrows show ‘triples’ (scale bar = 1 µm). (*iii*). Colocalization (white) of cb2 cell dendrites and GluR4 receptor subunit clusters. (*iv)* Cross-sectional view through a portion of the dendritic tree. Scale bar = 2 µm. **I.** Plot of contact fraction versus overlap volume for each innervated cone terminal in three cb2 and two cb3 cells. Dashed line indicates a contact fraction of 0.65.

To determine the fraction of presynaptic sites sampled by a cb2 cell, we plotted the peak amplitude of a bipolar cell epsc against the peak of the corresponding cone tpsc for a range of cone depolarizing steps including the responses shown in Fig 2B (green circles, Fig 2D). The resulting scatterplot was well-fitted by a straight line for cone responses <200 pA. Cone responses that exceeded this peak value were associated with a dramatic increase in the bipolar cell response amplitude up the maximum shown in Fig 2A. A rapid upswing in the bipolar cell versus cone response plot can occur if the cone transporter saturates during strong depolarizing stimuli while the lower affinity cb2 cell receptor (EC_50_ ∼350 µM (DeVries *et al*., 2006)) does not saturate as readily. Effective unitary event amplitudes were estimated from amplitude histograms (10.6 pA for the cone, Fig. 1F,G; 6.8 pA for the cb2). When responses were normalized by the unitary amplitudes, a straight-line fit (Fig. 2D) passed close to the origin (y-intercept = 0.04 quantal units; −0.24 ± 0.06 quanta, mean ± SE, n = 9, excluding one statistical outlier of −4.72 quanta), consistent with single event sensitivity in the bipolar cell. The linear fit had a slope of 0.648 suggesting that more 60% of the cone’s ribbon release sites were sampled by the cb2 cell’s dendrites. Over the linear region of the plot (Fig 2D), a fit to a power-law function, y = *a* + *b*x^c^, had an exponent, c, of 0.97 which is close to one (the linear model is preferred, p = 0.6239; F test). In 7 out of 10 cone to cb2 cell pairs, a linear fit was preferred over a power law fit. In three pairs, the power law fit was preferred (exponent = 1.72 ± 0.07). The average sampling of cone-released vesicles obtained by the scatterplot method was 47.9 ± 7.2% (mean ± S.E, n = 10; see Fig S2A-E for additional examples).

We used a second approach to determine the fraction of cone release sites sampled by a cb2 cell. In this approach, we assigned an effective quantal content to each tpsc irrespective of the cone voltage step during an experiment and then plotted the percentage of bipolar cell response failures as a function of quantal content (Fig 2F). An accurate count of the number of cone quanta depended on a well-defined cone response amplitude histogram, a criterion that was met for only 8 of 10 cone-to-cb2 cell pairs. Plots of percent response failure verses the number of cone vesicles released were fitted with ideal curves obtained by assuming 1) a fixed number of invaginating sampling sites relative to 20 ribbon active zones; 2) that invaginating dendrites respond to local release at the single quantal level; and 3) that the probability of release at each ribbon is the same (or nearly so) (Fig 2E). Using this approach, the cone to cb2 cell pair whose results were plotted in Fig 2D (Fig 2F, red circles) sampled 80% of the cone’s release sites. In the 8 experiments, contact numbers ranged from 4 to 16 out of 20 (40.6 ± 6.6% sampling; mean ± S.E.; Fig S2F). More generally, a mid-range slope of 1 when the data and model curves were plotted semi-logarithmically suggested that detection is fundamentally binary, with the fusion of one or more glutamate containing vesicles at a contact associated with success and the fusion of no vesicles with failure. For 6 of the 8 pairs, both analysis approaches produced comparable figures for the fraction of cone release sites sampled (Fig 2G).

To corroborate the physiological measurements of sampled release sites, we used STED microscopy to count the number of contacts between cb2 cell dendrites and overlying cone photoreceptors. Bipolar cells, including cb2 cells, were sparsely labeled with GFP by intravitreal injection of AAV (Fig 2H; supplemental movie 2). Individual cone terminals were marked by roughly circular groups of Ribeye and GluA4 subunits. The total number of ribbon release sites was estimated from close associations (using minimal surface to surface distance) between GluA4 and Ribeye (Fig 2H*ii*, open arrowhead; “doubles” in Fig 2I; see Methods for criteria). A subset of release sites (*i.e.*, doubles) was designated as contacted (Fig 2*ii*, arrows; Fig 2I, “triples”) based on both close association and volume overlap between GluA4 and dendritic labeling. In accordance with bipolar cell dendritic tiling (Wassle *et al*., 2009), we assumed that central cones in the dendritic field were solely innervated by the labeled cb2 cell (Fig 2H) and thus ordered the cone terminals according to contact fraction and focused on only the top quartile. The top quartile had a contact fraction of 80.7 ± 4.7% (n = 3 cb2 cells with 19.0 ± 4.4 presynaptic cones per cell) and corresponded to central cones. For comparison, a comparable analysis of labeled cb3 cell contacts (n = 2 cells) showed less volume overlap and a lower contact fraction (Fig 2I). Physiological measurements at the cone to cb2 cell synapse and STED-microscopic reconstructions of cb2 cell to cone contacts both support the idea that cb2 cells can sample most of a cone’s transmitter release sites.

### Sampling of the glutamate gradient by cb3 Off bipolar cells

We used the cell pair approach to characterize the cone sampling properties of cb3 cells. A sequence of depolarizing steps to −30 mV elicited cone tpscs that ranged up to −125 pA with a mean of −54.8 pA (Fig. 3A, upper left). The cone unitary event size equaled 21.7 pA when measured by amplitude histogram and 22.7 pA when calculated from P_0_. The postsynaptic cb3a cell (Fig. 3A, lower left) displayed a combination of responses and failures with a mean amplitude of −2.8 pA and an effective unit size of 3.1 pA (calculated from σ^2^/^-^*x*). The same voltage step during an earlier recording epoch produced a series of cone tpscs with a larger mean amplitude of −133.2 pA (Fig 3A, upper right). Associated bipolar cell responses (Fig 3A, lower right) had a mean amplitude of −33.9 pA and an effective unit size of 17.1 pA. The differences in effective unit size (*i.e*, 3.1 versus 17.1 pA) did not result from systematic differences in diffusion distance insofar as the time courses of the smaller and larger epscs in Fig. 3A were nearly identical (Fig S3A). A scatterplot of cb3a cell versus cone tpsc response (Fig 3B; Fig S3B-E) was fitted in the cone-linear range with a power-law function, y = Ax^b^, that had an exponent, *b*, of 1.72 (2.29 ± 0.09; n = 23 pairs; significantly different from 1.0; no difference between cb3a, 2.17 ± 0.52, n = 10, and cb3b, 2.45 ± 0.34, n = 12, cells; the axon of one cb3 cell was truncated precluding inclusion in a subcategory). Thus, the relationship between cone transmitter release and the bipolar cell response was non-linear.

**Figure 3.**
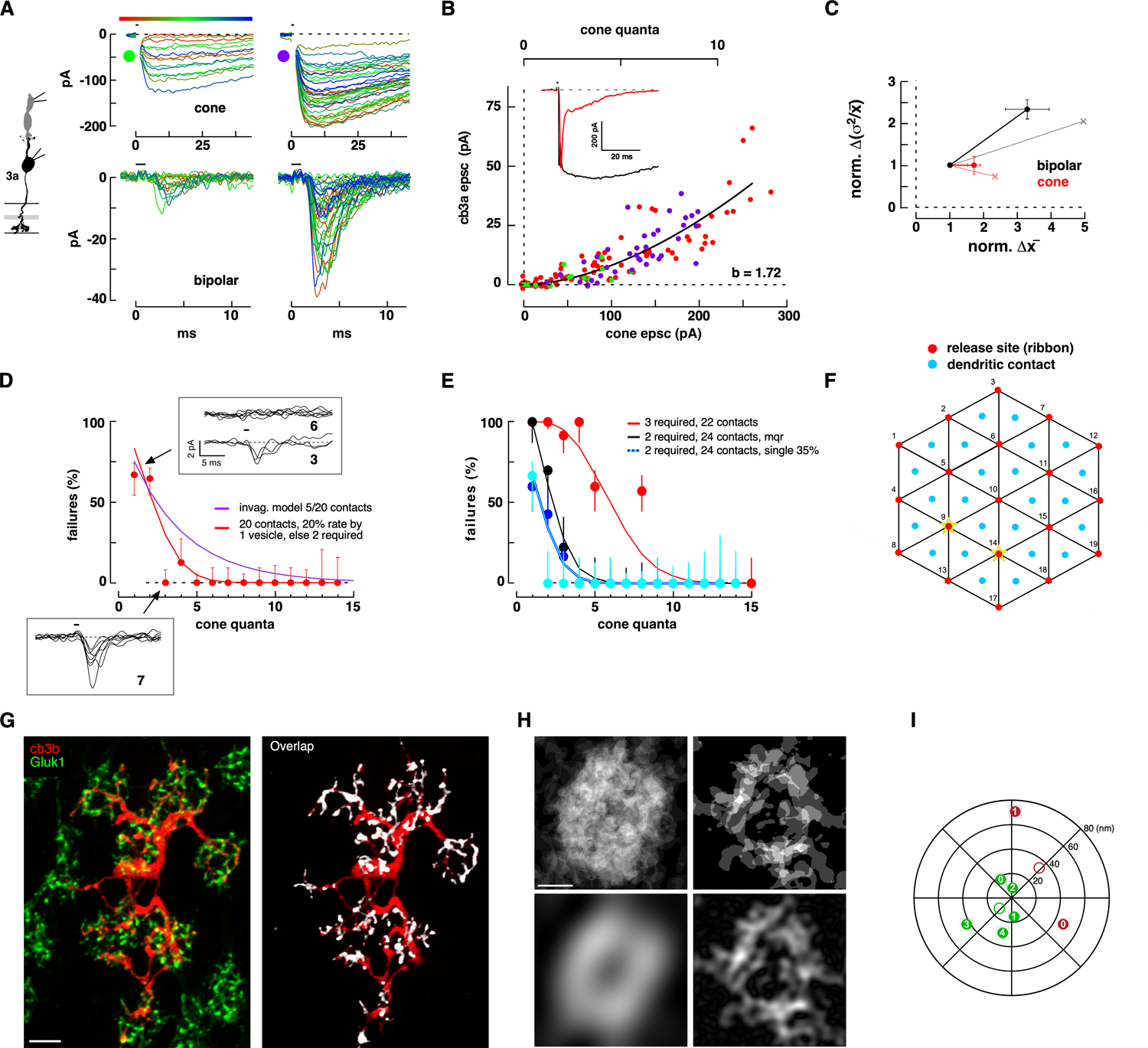
Non-linear responses at the cone to cb3 cell synapse. **A.** Paired responses obtained from a cone and cb3a bipolar cell during two stimulus epochs. **B.** Scatterplot of cb3a versus cone response. Green and magenta circles correspond to traces in A. *Inset.* Maximal cone tpsc (black trace) and bipolar cell epsc (red trace). **C.** Plot of fractional increase in effective unitary event size versus increase in the mean tpsc or epsc size for two epochs in each pair (n = 6 cone to cb3 cell pairs; mean ± S.E.). Normalization is to the smaller of the two response epochs (*e.g.*, A, left). Data from A is also plotted individually (crosses, light shading). **D.** Plot of percent bipolar cell response failures versus cone response quantal content for the cb3a cell in A,B. Insets show all individual responses (*i.e.*, successes and failures) at two representative cone quantal levels. An averaging procedure argued against the view that trials categorized as failures consistently contained small responses (Fig S3F). Theoretical curve from the model in Fig 2E with 5 out of 20 invaginations contacted (magenta). Red curve is from the model illustrated in F with a 20% detection rate at the single vesicle level, otherwise, a success occurs when an individual contact is exposed to two simultaneous release events. The model predicts that the bipolar cell samples cone release at 20 out of 24 potential contact sites. **E.** Percent failure plots for 4 additional cb3 pairs. Theoretical curves required a range of fit parameters including 0 – 35% effectiveness at the single vesicle level, a dendritic contact threshold of 2-3 simultaneously release vesicles, and either 0 or 20% of the events were multiquantal (mqr). The model predicted occupancy at 22 – 24 contact sites. **F.** Diagram of the model showing 19 ribbon release sites (red circles), two of which are active (yellow stars), and the locations of 24 potential dendritic contact sites. **G.** (*left*) STED microscopic image of a retinal whole mount with labeled GluK1 subunit (green) and a cb3b cell (red). (right) Colocalization of GluK1 and cb3b labeling (Scale = 2 µm). **H.** (*left, upper*) GluK1 labeling from individual cone terminals from the entire image a portion of which is shown in G (*left*) were binarized, superimposed, and point-by-point averaged. (*left, lower*) The profile was smoothed (see Fig S4A,B for procedural details and additional examples). (*right*) Averaged binarized profiles of the cb3b cell dendrites beneath six of the cones with largest amount of colocalization (Fig S4C). Scale = 1 µm. **I**. Polar plot of weighted centroid displacement obtained from five images with GluK1 labeling (filled green circles, the open circle is the average; Fig S3B). Centroid locations for two of the images that contained cb3 cells (red circles; Fig S3C).

As bipolar cell responses increased in amplitude, the effective unitary event size also increased (Fig 3A). We tested whether the increase in the effective bipolar cell unit was associated with a comparable increase in effective unit size in the cone response, as might occur if there was an increase in the proportion of multiquantal events. In this test, we used σ^2^/^-^x as a qualitative measure of response amplitude, both for cone tpscs and bipolar cell epscs, and calculated the relative change between stimulation epochs in the same recording (*e.g.*, Fig 3A, left and right). The larger amplitude epoch was selected to have cone responses within the linear summation range (121.8 ± 33.7 pA, mean ± S.D.; n = 6 cone to cb3 cell pairs). Between epochs, cone responses increased by a factor of 1.71 ± 0.19 (mean ± S.E.; p = 0.0135 one sample t-test, Fig 3C) while bipolar cell responses increased by a factor of 3.29 ± 0.65 (p = 0.0169 one sample t-test; ratio difference is significant, p = 0.156, Wilcoxon signed test for paired data). The 1.71-fold increase in the cone response occurred in the absence of a change in the variance to mean ratio (1.0 ± 0.21) suggesting an underlying increase in the quantal content of the average tpsc without a change in effective unit amplitude. In contrast, the 3.29-fold increase in mean bipolar cell epsc amplitude was accompanied by a 2.33 ± 0.23-fold increase in the variance to mean ratio (p = 0.0022), suggesting that an increase in the mean response was associated with both an increase in unit size and quantal content. The implication is that the size of the effective unitary event near threshold, as reported by cb3 cells, increases with mean cb3 bipolar cell response amplitude.

We next characterized the sampling properties of cb3 cell dendrites by plotting percent response failure versus the number of cone quanta released. A postsynaptic cb3a bipolar cell responded ∼30% of the time when the presynaptic cone released a single quantum of transmitter (Fig 3D). Two additional cb3 cells had a similar percent response at the single vesicle level, while two cb3 cells in the sample did not respond at the single vesicle level (Fig 3E). Reponses transitioned to a 90-100% success rate when cone tpscs contained 4-5 vesicles (4 out of 5 pairs, Figs 3D,E). Plots of percent response failure versus cone quanta released were poorly fitted by the model used for cb2 cells (Fig 3D, magenta curve). Rather, response plots were fitted by a model in which a “success” depended on the exposure of any of the cb3 cell cone contacts to two or more overlapping release events from up to three neighboring invaginations (Fig 3D, red curve; Fig 3E; total model contacts = 24). This type of model could fit the rapid transition from a complete response failure when one vesicle was released by a cone to no failures when 4-5 vesicles were released (e*.g.*, Fig 3E, black curve). To account for the observed rate of failures at the single vesicle level, the model could be adjusted so that a variable percentage (*e.g.*, 30%) of individual vesicles elicited a bipolar cell response (Fig 3D, red curve; Fig 3E, cyan and blue curves). In one of the five pairs, it was necessary to assume a cooperativity of 3 events (Fig 3E, red curve). In all cases, the model suggested that an individual cb3 bipolar cell sampled release from nearly all of a cone’s invaginations (range = 80-100%) with a threshold that excluded at least 65% of the individual release events.

We used STED microscopy to examine the contact patterns of cb3 cells beneath individual cone terminals. Both cb3a and cb3b cells express kainate receptors containing GluK1 subunits, and the volume beneath each ground squirrel cone terminal contained a 3-dimensional web of GluK1 labeling (Fig 3G, left; see supplemental movie 3). We analyzed the pattern of GluK1 labeling at the cone terminal by binarizing a collapsed stack from a 3D image, fitting an identical square around each terminal, and averaging the images across all squares to calculate the sampling profile and mean density (Fig 3H, upper left; Fig S4A). We summed image squares without rotation on the assumption that radial differences, if they existed, would have the same relative orientation (*e.g.*, toward the optic nerve head) in a small retinal region. Smoothed profiles (Fig 3H, lower left; Fig S4B) consisted of a circular or elliptical ring with an average density of 71.1 ± 9.3% (mean ± S.D., n = 5; Table S1) and a central a dimple with a density of 63.0 ± 13.2%. A weighted vector sum of the density within the ring had X-Y coordinates very close to the unweighted mean (Fig 3I, green circles; Fig S4B; Table S2) consistent with a uniform radial density. The dendrites of GFP-filled cb3 cells (Fig 3G, right; Fig S4C; n = 2 of the 5 analyzed images) colocalized with GluK1-labeled profiles beneath a cone. Inspection suggested that major dendrites gave rise to branches that sample from all regions of cone terminals (Fig 3G, right). A qualitative analysis of cb3 dendritic profile overlap beneath cones (Fig 3H, right, I; Fig S4C) provided additional confirmation for a ring-like distribution. Since the dendrites of bipolar cells within a type tile the retina with minimal overlap (Wassle *et al*., 2009), each cone should receive innervation from the equivalent of one cb3a and one cb3b cell. This dual innervation accounts for nearly all GluK1 labeling insofar as cb1 cells make relatively few cone contacts and have weak GluK1 labeling (see below). The high density of GluK1 labeling beneath a cone terminal combined with the ring-like coverage of cb3 cell dendrites supports the idea of comprehensive sampling of a cone’s transmitter release sites.

### Sampling by cb1a Off bipolar cells

We next characterized the sampling properties of cb1a cells and observed a marked non-linearity. In a cb1a cell pair, a train of brief cone depolarizations, first to −40 and then to −35 mV produced a series of cone responses with discrete levels (Fig 4A, upper panel). Of the 25 trials that resulted in a cone tpsc, only the largest tpsc, corresponding to a content of at least 7 quanta, resulted in a cb1a cell epsc (Fig 4A, lower panel, thick lines). A scatterplot (Fig 4B) showed that the cb1a cell response did not deviate from baseline until the cone response amplitude exceeded ∼125 pA, equivalent to 5 cone events (Fig 4C). Between cone response amplitudes of 125 and 300 pA, the bipolar cell response slowly increased, and then increased more rapidly when cone responses exceeded 300 pA, a consequence of transporter saturation. The initial portion of the scatterplot, within a cone-linear range of 10 vesicles, was fitted with a power-law curve that had an exponent of 1.80 (3.23 ± 0.48, n = 8 cone to cb1a cell pairs and 2.00 ± 0.12, n = 2 cb1b cell pairs, not significantly different p = 0.272 for a total of 3.03 ± 0.41, n = 10; different from cb3 cells, p = 0.0188; see Fig S5A-C for additional scatterplots). We used a second approach to characterize response non-linearity that allowed us to add two cone-cb1a cell pairs to the sample where the deviation of the cb1a cell response from baseline was negligible over the entire cone-linear range. On the limiting idea that a shallow slope might result from a single dendritic contact linearly sampling a single cone invagination (Fig 1C), we fitted the scatterplot with a line over a cone tpsc range equivalent to 7 quanta (Fig 4B, solid cyan line). From the slope, we calculated the cb1a cell epsc current increment (0.34 pA) per cone tpsc event (25.1 pA), and then extrapolated how many increments would be required to produce the maximal cb1a cell response if summation were linear (dashed cyan line). Applying this approach, we calculated that the maximal epsc contained at least 548 quanta (median = 1122 quanta per maximal epsc; range = 167 to infinity; n = 10). For comparison, a maximal 1 ms stimulus caused the fusion of ∼20 vesicles from an individual ribbon site or ∼360 vesicles from the entire terminal pool (see below). Thus, cb1a responses increase supra-linearly with the amount of cone transmitter released.

**Figure 4.**
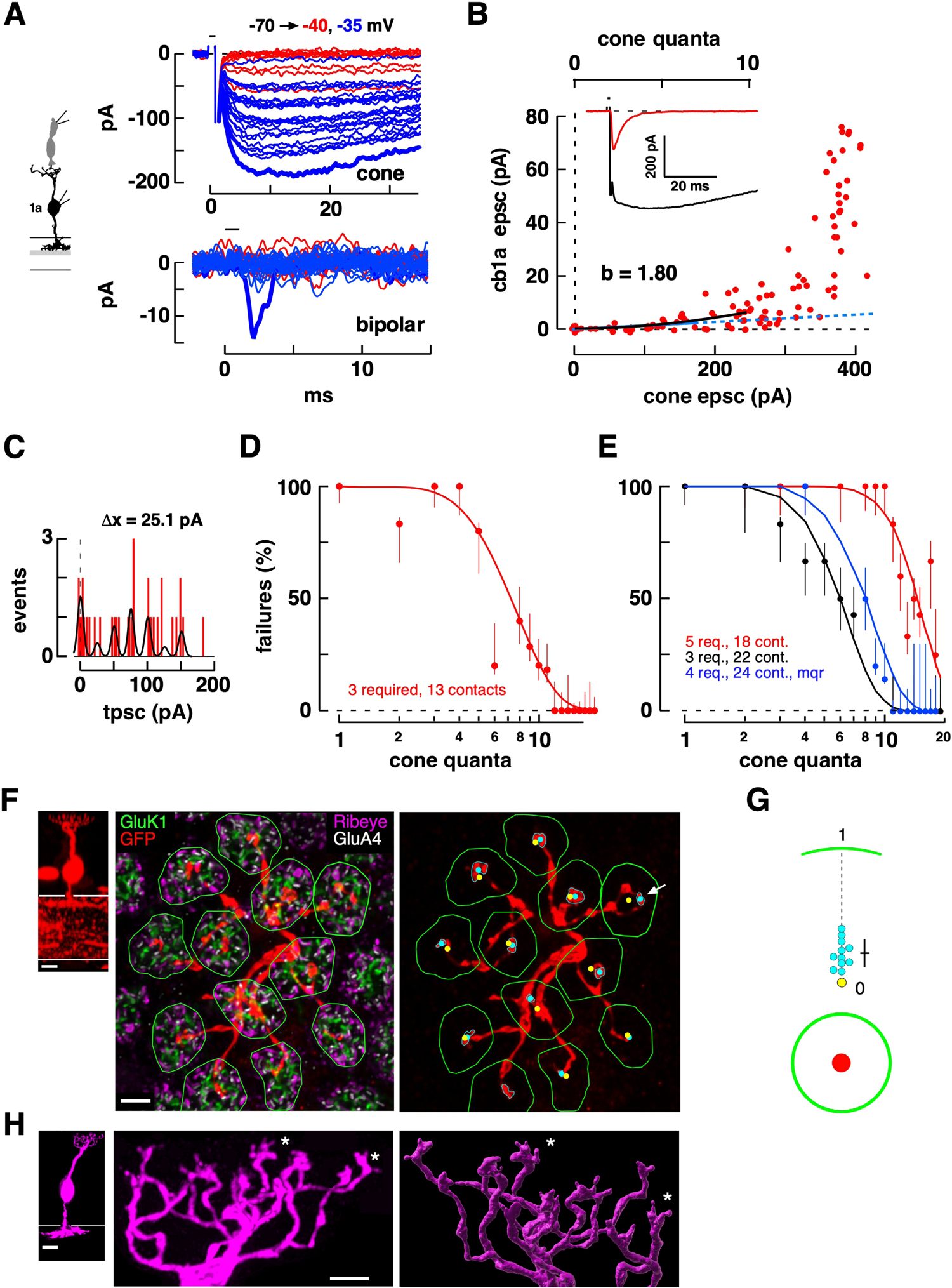
Non-linear responses at the cone to cb1a cell synapse. **A.** Pre- and postsynaptic responses in a cone to cb1a cell pair. Sequential responses to steps to −40 and −35 mV are superimposed. **B.** Scatterplot of cb1a peak epsc versus cone tpsc responses during several epochs including those shown in A. The black curve is a fit to a power law fit. The solid and dashed cyan lines are a linear fit and its extrapolation, respectively. *Inset:* Maximal cone and bipolar cell responses. **C.** The amplitude histogram from the cone responses in A were fitted with a multipeak Gaussian. The calculated unitary event amplitude was 25.1 pA. **D.** Plot of percent response failures versus cone tpsc quantal content. **E.** Percent response failure plots for 3 additional cb1a cells. Best fits were obtained by assuming that 3-5 vesicles were required to produce a success at a contact, with sampling of 18 – 24 out of 24 ribbon release sites. **F.** (left) Confocal image of a GFP labeled cb1a cell (red; scale = 5 µm). (middle) Maximum intensity projection of a 3D STED image of the cb1a cell’s dendrites (red) in the wholemount configuration. Terminals were labeled for GluK1, GluA4, and Ribeye (scale = 2 µm). Outline of each contacted cone terminal is shown (green polygons). (right) Image of cb1a bipolar cell dendrites (delimited by cyan polygons) with superimposed polygonal outlines for the cone terminals. Centroids were calculated for each contacted cone terminal (yellow dot corresponds to green polygon) and each dendritic terminal profile (cyan dot). **G**. (upper) The effective radius of each terminal region was calculated from a circle of the same area and used to normalize the distances of the contact centroid from the terminal centroid (mean ± S.D. normalized micron distance = 0.219 ± 0.104; significantly different from 0, p = 0.0001). (lower) Area ratios for the contact versus terminal (2.92 ± 2.1%). **H.** A different cb1a cell showing a collapsed stack of STED images obtained in the slice orientation. A surface fit to the dendritic tree shows terminal bulbs with up to 5 thin extensions. Asterisks indicate the same terminal dendrites viewed from slightly different orientations. Scale bars (10 µm, left; 2 µm center and right).

We next characterized the sampling properties of cb1a cell dendrites by plotting percent response failures versus the number of cone quanta released (Fig 4D,E; Fig S5). Response failures were typically 90-100% in the range of 1-4 vesicles, transitioning to 0% failure rate when 11 or more quanta were released. Applying the same model used for cb3 cell responses (Fig 3F), cb1a cell responses required the local overlap of 3-5 events at a contact while sampling release from between one-half and nearly all of a cone’s release sites. Models with less cooperativity including a linear summation model or one depending on the summation of only two events could not account for the steep transition between failure rates of 100% and 0%.

In reconstructions (Fig 4F), cb1a cell dendrites had a prominent bulb-like expansion that was confined to the central region of the cone terminal (Fig 4G; Fig S5E-H; supplemental movie 4). Remarkably, central targeting was maintained even when dendrites divided into separate branches well before reaching a terminal (Fig 4F, arrow). High resolution images of the dendritic bulbs (Fig 4H) showed that the terminal expansion ended in 2-5 short, fine processes that extended further toward the cone. Due to a weak association between GluK1 labeling and cb1 cell dendrites, we could not determine whether postsynaptic receptors were localized to the bulb or fine processes. Thus, cb1 responses to cone transmitter release were highly non-linear, and cb1a cells made only a few dendritic contacts in the central region of the cone terminal.

### Simultaneous epscs in Off bipolar cells

In the cone to cb1a cell scatterplots, the non-linear relationship is obscured by the onset of cone transporter saturation. To better visualize the synaptic non-linearity, we took advantage of the linear relationship between cone transmitter release and the cb2 epsc response, which we assume continues beyond the transmitter levels at which cone transporters saturate. Next, we further characterized the cb1a cell non-linearity by evoking transmitter release from a presynaptic cone and measuring the simultaneous responses in postsynaptic cb1a and cb2 cells (Fig 5A-C). In both cases, the cone was depolarized for 1 ms in the loose seal mode. Loose-seal recording provides both stable epsc responses over long time periods (Lindstrom *et al*., 2014) and allowed us to find the relatively rare cones that are presynaptic to both bipolar cells through a trial-and-error process. The scatterplot of cb2 versus cb1a peak epsc response had at least two regions: At low cone stimulus strengths, cb2 cells responded but cb1a cells did not; at medium cone stimulus strengths, the responses of both cells were linearly related; and finally, during the strongest pulses, cb2 cell responses may saturate (Fig 5B; (Grabner *et al*., 2016)). Fits to the linear portions of the plots in Figs 5B,C had y-intercepts equivalent to about 28 and 9 cb2 cell quanta, respectively. For comparison, scatterplots of responses from cb3b and cb3a cells each paired with cb2 cells (Fig 5D,E) displayed linear relationships with intercepts closer to the origin (*i.e.*, both ∼3 quanta). Thus, consistent with the results from cone to bipolar cell pairs, the results from paired bipolar cells support a substantial threshold non-linearity in cb1a cells and a lesser threshold non-linearity in cb3 cells. Notably, response differences do not result from sampling different pools of presynaptic ribbons as correlated fluctuations in the trial-to-trail responses of the postsynaptic pairs during an epoch (Fig S6) imply that cb1, cb2, and cb3 cells sample from overlapping ribbon populations.

**Figure 5.**
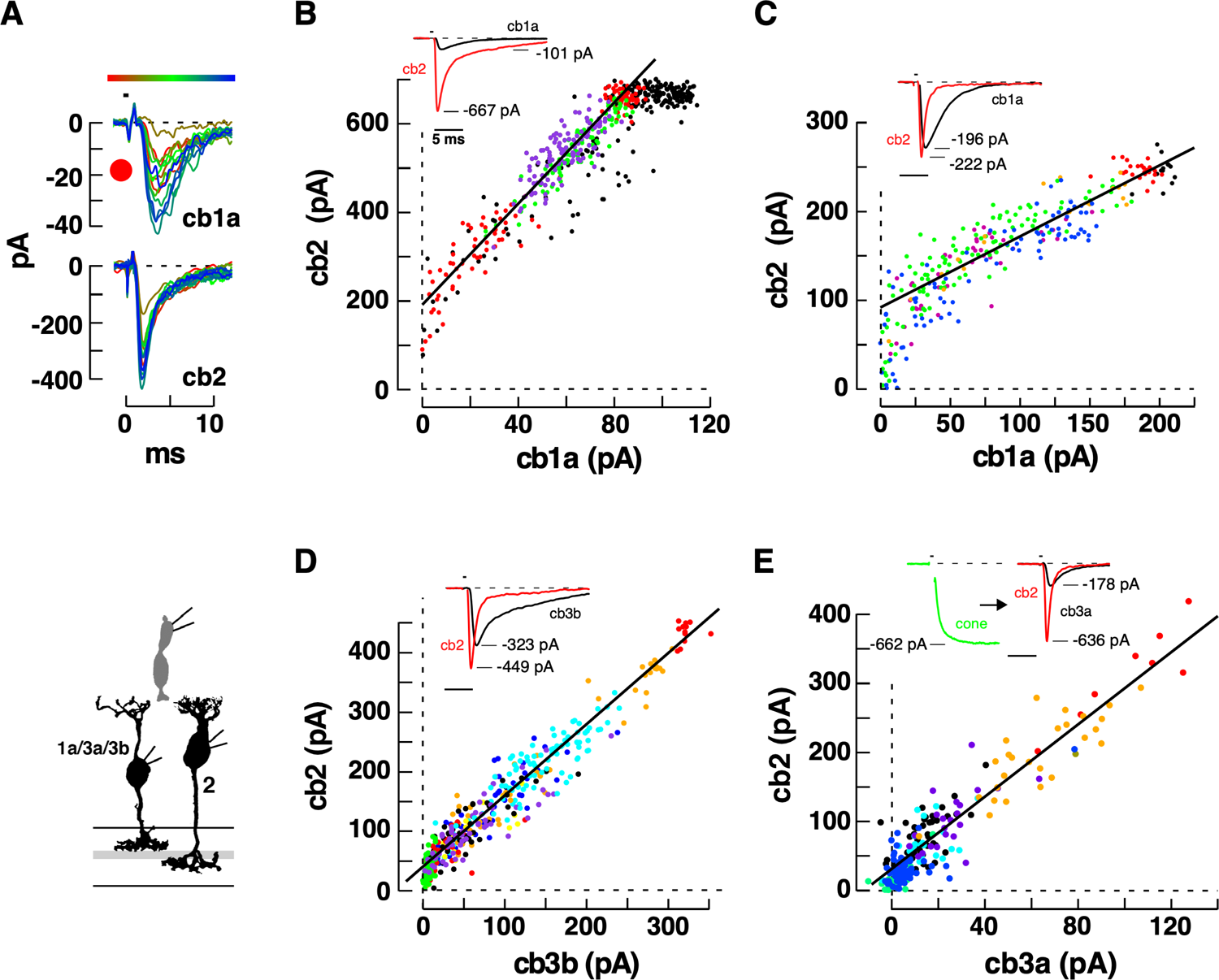
Simultaneous responses measured from two bipolar cells connected to a single stimulated cone. **A.** (upper) Paired epscs for a cb1a and cb2 cell during a train of 1 ms pulses in the presynaptic cone applied in the loose seal configuration. (lower) Diagram of recording configuration. **B.** Scatterplot of peak cb2 versus peak cb1a epsc response obtained during several epochs (linear fit: slope = 5.7 and y-intercept = 190 pA). Epochs shown in different colors except when pulses to two different voltages alternated between trials. The traces in A correspond to the lower set of red points. Linear fit excludes the highest and lowest points. **C.** Scatterplot for a second cb1a and cb2 cell pair (slope = 0.8; y-intercept = 90 pA). **D.** Scatterplot for a cb2 and cb3b cell pair (slope = 1.2 and y-intercept = 40 pA). **E.** Scatterplot for cb2 and cb3a cell pair (slope = 2.6 and y-intercept = 31 pA). Insets show the peak epsc responses for B-D and a triple whole cell recording for E.

### Differences in glutamate affinity between cb1 and cb3 cell receptors

To determine the basis for the distinctive threshold in cb1 cells, we compared the concentration-response properties of cb1a and cb3 cell receptors by removing bipolar cell somas and exposing them to rapid step changes in glutamate concentration (Fig 6A,B). AMPA receptors, which comprise ∼20% of the postsynaptic pool in cb3 cells (Lindstrom *et al*., 2014), were blocked with GYKI53655 (35 µM). By preserving the tracer-filled axon and connected dendrite in the slice, the bipolar cell type could be unambiguously identified after recording (Fig 6B, right). Somatic cb3 and cb1a responses to an 18 mM glutamate step had peak amplitudes of 250.6 ± 73.4 (mean ± S.D., n = 7) and 40.9 ± 20.6 pA (n = 6), and 20-80% rise times of 0.236 ± 0.136 and 0.192 ± 0.061 ms, respectively. A plot of glutamate concentration versus peak response for the cb3a cell in Fig 6B (upper panels) had an EC_50_ of 0.53 mM (0.37 ± 0.06 mM; n = 7; mean ± S.E.; Fig 6C,D) and a Hill coefficient of 0.77 (1.09 ± 0.17) (Fig 6D). The cb1a cell in Fig. 6B (lower panels) had an EC_50_ of 1.57 mM (1.40 ± 0.16 mM; n = 6) and a Hill coefficient of 1.2 (1.11 ± 0.08; not significantly different cb1 versus cb3, p = 0.922) for an overall 3.82-fold difference in EC_50_ (cb1 versus cb3, p < 0.0001). In a separate set of rapid perfusion experiments, we measured the responses of cb1a and cb3 cell somatic receptors to glutamate concentrations between 12.5 and 100 µM, a range that modeling (see below) suggests encompasses the concentration of glutamate at cone basal contacts following single vesicle fusion (Fig S7A-D). A 5-10-fold difference in glutamate sensitivity was maintained in this low concentration range. Receptors also differed with respect to IC_50_ (3.66-fold higher in cb1a versus cb3 cells; Fig S7E-H) and recovery from desensitization (faster in cb1a cells; Fig S7I,J) consistent with a previous report (DeVries, 2001).

**Figure 6.**
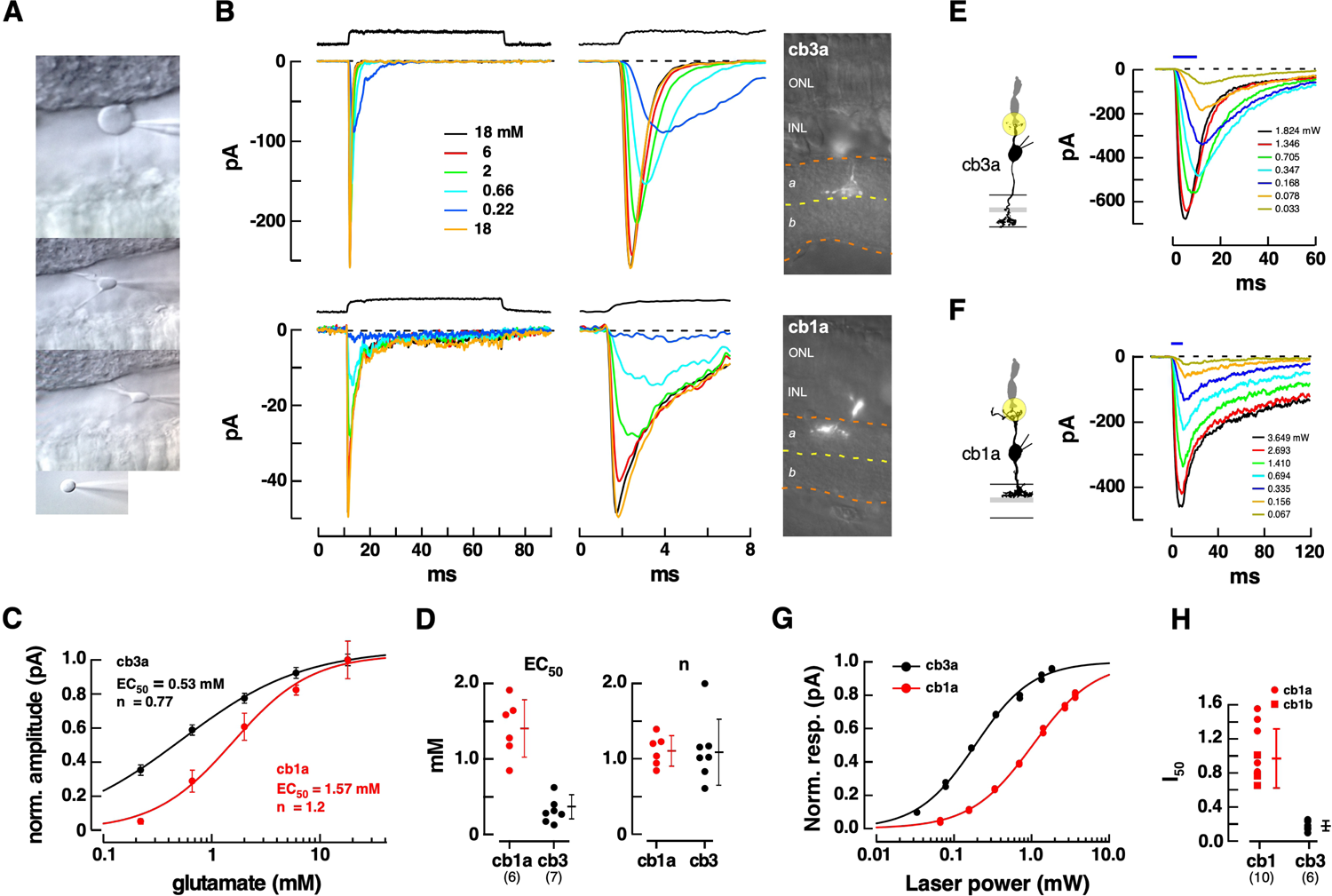
Responses of isolated cb3 and cb1 cell kainate receptors to concentrations of glutamate. **A.** Sequence illustrates the procedure for establishing a whole cell recording from a bipolar cell and withdrawing its soma from the retina while leaving an intact axon and dendrite in the slice. **B.** Responses of somas to rapid applications of glutamate (0.22 − 18 mM). Solution exchange temporal profiles (black traces, above) were obtained from the open-tipped pipette following cell rupture. The entire step response is shown on a slow time base (left) and the peak response on a faster time base (middle). Fluorescence images of the remaining axon/dendrite allow identification of the cell type (right). **C.** Plot of normalized peak response versus concentration for the cells in B. Points are the average of several repeats (mean ± S.D.). **D.** Categorical scatterplots of EC_50_ and ‘n’ obtained from the Hill fits for cb1a and cb3 cells showing the mean ± S.D. Number of cells in parentheses. **E., F.** Responses of a cb3a and cb1a cell to glutamate uncaging. MNI-glutamate (5 mM) was applied by puffer to the OPL and a 3 µm diameter uncaging spot, illustrated by the yellow circle, was focused on at the dendrites of the recorded bipolar cell at the cone terminal. The amount of released glutamate was controlled by adjusting flash intensity (mW; flash duration shown above). 20-80% rise times were 1.23 and 2.17 ms for the maximal cb3a and cb1a responses, respectively. **G.** Plots of normalized peak response versus spot intensity for the cells in E and F. Responses were fitted with a Hill Equation to obtain I_50_. **H.** Plots of I_50_ for individual cb1 and cb3 cells showing the mean±S.D. AMPA receptors were blocked with 35 µM GYKI 53655 in all experiments.

To address the possibility that dendritic and somatic kainate receptors could have different EC_50_s, we measured the difference in receptor sensitivity between cb1 and cb3 cells during laser-induced glutamate uncaging. MNI-glutamate (mM) was uncaged using a 3 µm diameter, variable intensity spot aimed at the synaptic contacts between a cone and a recorded bipolar cell (Fig 6E,F). cb3 cell responses had an I_50_ (I, laser intensity) of 0.18 ± 0.02 mW. (mean ± S.E.; n = 6 cones), 5.8-fold lower than the I_50_ of cb1a cells (1.04 ± 0.17 mW; n = 7; significantly different, p = 0.0007). The results suggest that differences in receptor affinity can play a role in the different synaptic responses of cb1 versus cb3 bipolar cells.

### Role of basal glutamate binding sites

We next considered whether the glutamate transporters at the cone terminal might function as saturable binding sites that reduce the amount of transmitter that can reach cb1 and cb3 cell contacts (Rowan et al., 2010). The size of the maximal glutamate transporter current, typically 500 – 800 pA at −70 mV (Szmajda and Devries, 2011), combined with published values for single channel conductance (0.45 pA at −70 mV; (Schneider et al., 2014)) suggest that a cone terminal may contain at least 20,000 transporters (see (Hasegawa et al., 2006), for a higher density estimate at the rod terminal). We tested for a role of transporters in glutamate binding and response attenuation at the quantal level by applying TBOA and measuring the change in postsynaptic response amplitude and time course. Cone transmitter release was evoked by a train of depolarizing pulses while simultaneously recording cone tpscs and bipolar cell epscs. In a cb3a bipolar cell (Fig 7A,B), TBOA completely blocked the cone response while producing a 3.48-fold increase in epsc amplitude (4.23 ± 1.05-fold for n = 11 cb3 cells; Fig 7B,E), a 2.80-fold prolongation of the epsc response decay (2.39 ± 0.16-fold, n = 11; p < 0.0001; Fig 7B), and an −7.56 pA increase in bipolar cell baseline current (−7.70 ± 2.00 pA, n = 13; p = 0.0023, different from no change). The amplitude of the epsc at a cone to cb1a cell synapse was enhanced by 3.53-fold by TBOA (4.21 ± 1.29-fold; Fig 7C,D). Response decay was prolonged by 1.44-fold (1.49 ± 0.94-fold, n = 10; p = 0.0005) and baseline current was elevated by −0.38 pA (−9.02 ± 3.46 pA, n = 11; p = 0.0263). Increases were observed under control conditions that allowed recordings at cone to cb2 cell synapses, which demonstrated no change in epsc amplitude (Fig 7E), and in 35 µM GYKI53655, which prevented horizontal cells from depolarizing in response to elevated concentrations of glutamate and possibly modulating cone transmitter release (Jackman et al., 2011; Wang et al., 2014). With GYKI53655 present, it was not possible to record cb2 cell responses. The epsc increases in cb1 and cb3 cells were similar whether measured in GYKI53655 (n = 4 for cb1 and 3 for cb3 cells) or in control (p = 0.3843 and 0.7863, respectively; Fig 7E).

**Figure 7.**
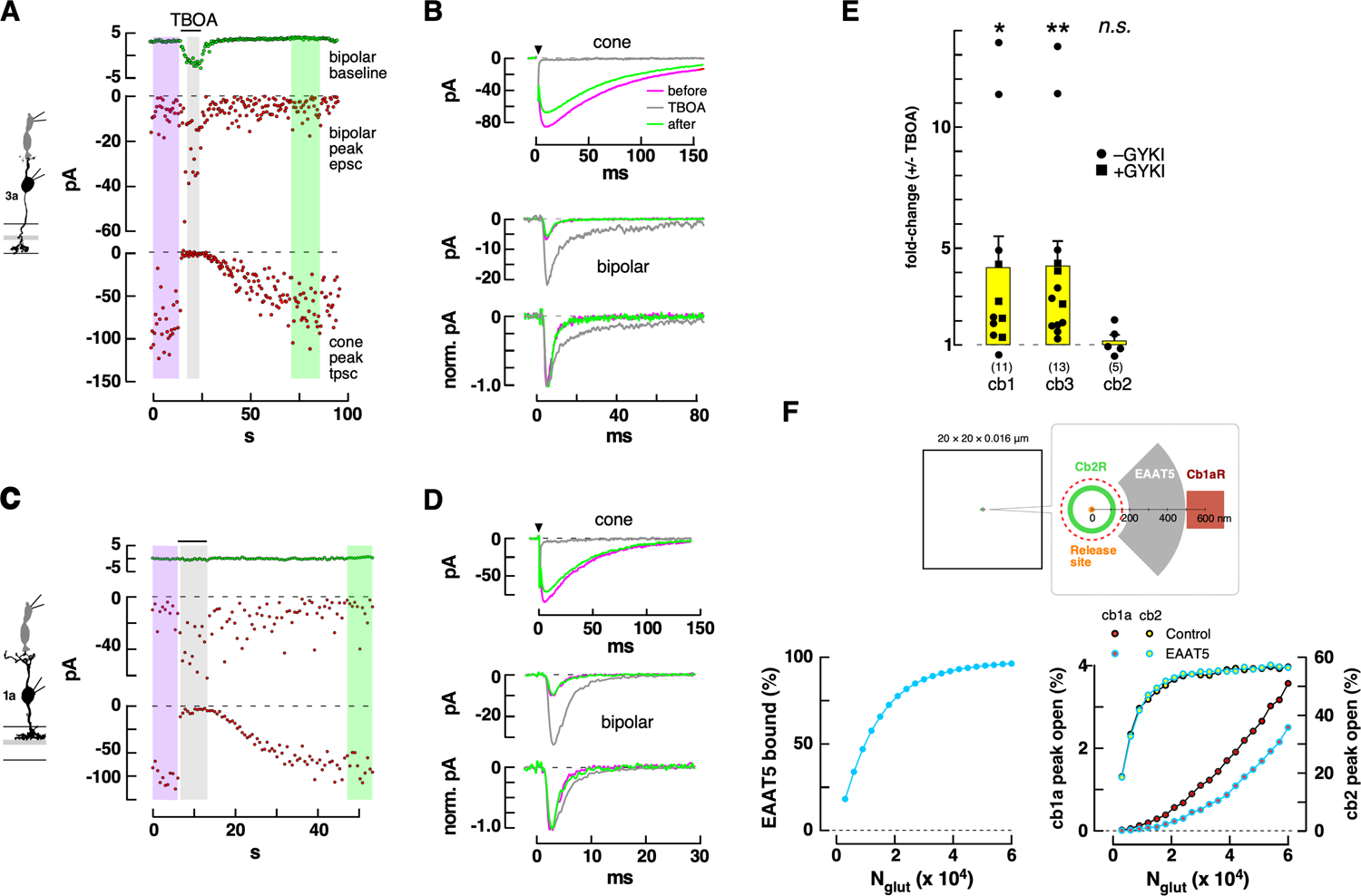
Effect of TBOA (325 µM) epsc amplitude in Off bipolar cells. **A.** Plots of bipolar cell baseline current (top), epsc amplitude (middle), and peak cone transporter current (below) as a function of time during a train of 1 ms steps from −70 to −30 mV. **B.** Average responses before, during, and after puffer application of TBOA. Color coding corresponds to the shading in A. **C,D.** Similar results for a cone to cb1a cell pair. Step from −70 to −30 mV. **E.** Aggregate results (*, 0.05 > p > 0.01; **, 0.01 > p > 0.001). Experiments on cb1 and cb3 cells were performed both with and without the addition of GYKI53655. Results in A,B were obtained without and in C,D with GYKI53655. **F.** Monte Carlo simulation of the effect of cone EAAT5 transporters on cb1a and cb2 cell receptor responses to released glutamate. Diagram of the model cleft (above). (lower, left) Peak EAAT5 glutamate binding as a function of the number of released glutamate molecules (N_glut_) for 3200 EAAT5 transporters in a 0.165 µm^2^ patch (gray area). (lower right) Peak channel open probability (percent) for an annulus containing 100 cb2 cell AMPA receptors (green) or a patch containing 350 cb1a cell receptors (0.04 µm^2^). Simulations on each receptor were performed separately while varying the number of transporters. Red dashed circle delimits the extent of an invagination in this model.

We next wanted to determine whether basally located transporters could contribute to the non-linear responses in cb1a cells. To assess the impact of basal transporter localization on Off bipolar cell responses, we created a simplified Monte Carlo model of glutamate diffusion from a release site into the cone synaptic cleft. The model assumed that transporters were localized to patches of basal membrane and acted as reversible binding sites on the time scale of an epsc. We found that modeled cb1a cell receptor responses to glutamate were moderately non-linear over the tested range even in the absence of transporter (*i.e.*, under control conditions). At low glutamate concentrations, transporters located between release sites and cb1a cell postsynaptic receptors could further reduce the amount of glutamate that reached receptors and nearly abolished responses (Fig 7F; Fig S8). At higher glutamate concentrations, transporter binding sites saturated (Fig 7F, right) and the effect on receptor response was relatively reduced as signified by the upward curvature of the cb1a cell receptor response and its subsequent parallel rise relative to control (Fig 7F). We also observed that adding transporters to the opposite side of the release site relative to a receptor patch or beyond the receptor patch (*e.g.*, on distant Muller cell processes) had little effect on peak epsc responses in the model as illustrated by the early linear rise of the cb2 cell receptor response. The results suggest that saturable sites on glutamate transporters can bind and buffer glutamate at low levels of transmitter release, reducing the activation of the receptors on postsynaptic cb1 cells and potentially contributing to a non-linear response. A lesser effect of transporter blockers on cb2 cell epscs, and therefore a more linear relationship between glutamate release and receptor response, can be understood if transporters are located outside of the invaginating contact sites (Fig 7F).

### Spatiotemporal glutamate gradient at the cone terminal

Cone photoreceptors rest at a relatively depolarized voltage in the dark where they undergo a steady rate of vesicle fusion. If the rate is relatively low, fusion events at the ∼20 ribbons in a cone will effectively be spatiotemporally discrete, each resembling those evoked by a small 1 ms depolarization. The properties of the steady response can then be inferred from those of the small evoked responses. Alternatively, a high maintained rate of fusion may lead to a spatially uniform gradient at the terminal, dissimilar from that produced by a brief cone depolarization.

We first established the ribbon as the primary site of vesicle fusion during a brief depolarizing pulse. The number of vesicles in the membrane-docked pool was determined by measuring the change in cone whole-cell membrane capacitance (ΔC_m_) during a series of depolarizing steps from −70 to −10 mV (Fig. 8A, a step to −10 mV produced a maximal capacitance change; data not shown). A 1 ms step caused a ΔC_m_ of ∼16 fF, while longer steps up to 30 ms led to incremental gains (Fig. 8B). A plot of average ΔC_m_ versus step duration was best fit with a bi-exponential function that gave a time constant (τ) for the first component of fusion equal to 0.4 ± 0.2 ms and a τ_2_ = 7.3 ± 3.8 ms, with amplitudes of 15.6 ± 3 and 10.5 ± 2.5 fF, respectively (n = 7 cones; Fig 8B). We designated these two rapid kinetic phases as ultrafast and fast, respectively, with reference to comparable pools in goldfish Mb1 bipolar cell terminals (Grabner and Zenisek, 2013). Based on a mean single vesicle capacitance of 43 attofarads (aF, 10^-18^ F; Fig S8K; specific membrane capacitance = 0.9 µF-cm^2^, (Grabner and Moser, 2018)), the ultrafast and fast pools contained ∼360 and ∼240 synaptic vesicles (SVs), respectively, together creating a readily releasable pool (RRP) of 600 SVs.

**Figure 8.**
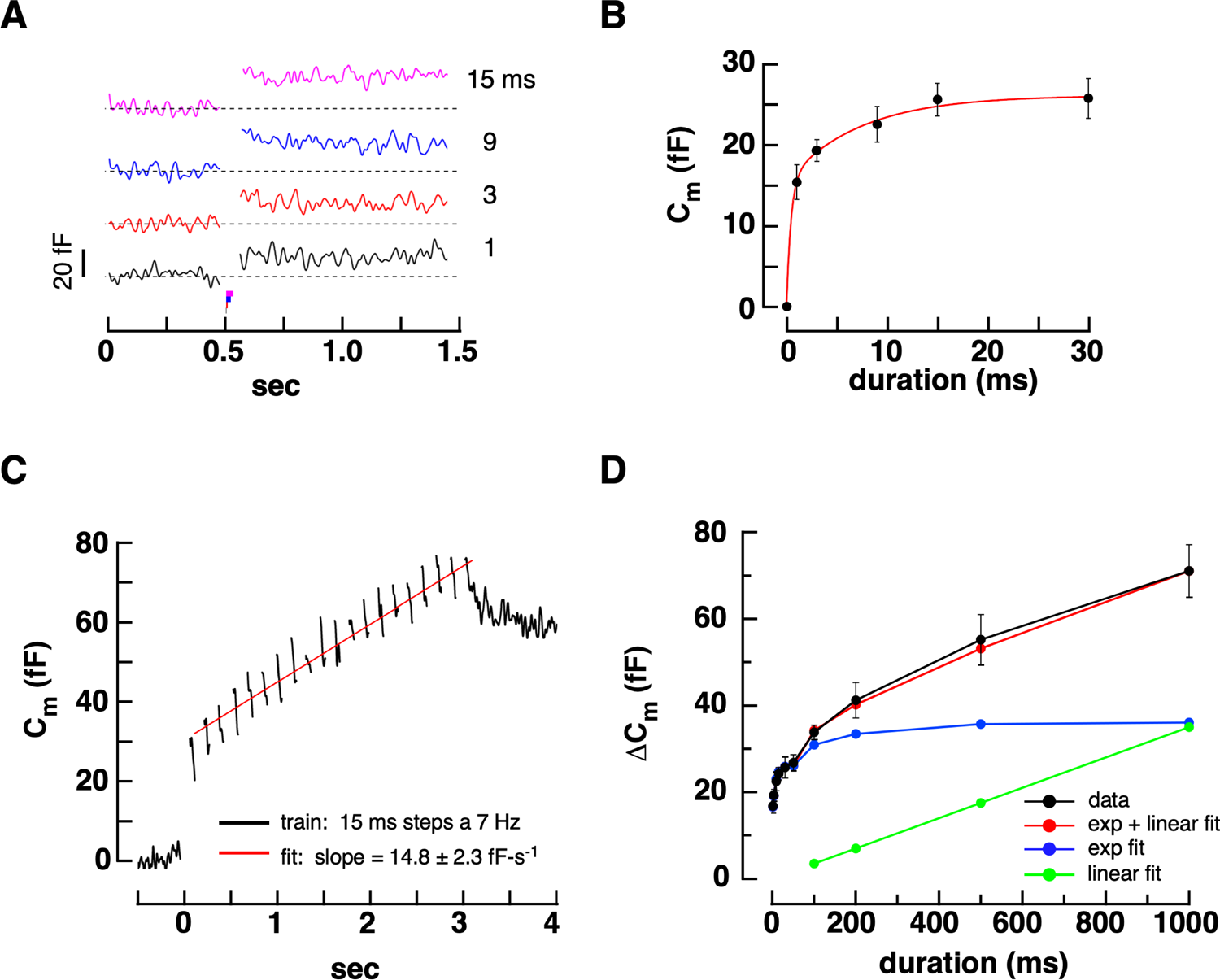
The steady rate of vesicle fusion at the cone terminal. **A.** Increases in membrane capacitance produced by stepping cone membrane voltage from −70 to −10 mV for durations of 1-15 ms. **B.** Plot of ΔC_m_ versus step duration fitted with a bi-exponential curve (r = 0.98). **C.** Increase in C_m_ during a train of 20 x 15 ms steps to −10 mV at 7 Hz. Average of 8 cells. Intercept = 30.6 fF. **D.** Plot of average ΔC_m_ versus step duration (−70 to −10 mV; black circles and straight lines; 9 cells; mean ± SE). The plot was approximated by summing two components, a bi-exponential fit to the rise at early times (blue circles and lines) and a linear fit to the ΔC_m_ values between 0.2 and 1.0 s (green points and line). The steady replenishment rate is obtained from the slope of the green line.

We next compared the number of vesicles in the RRP to the number of ribbon and membrane associated vesicles from EM reconstructions of the cone terminal (Fig S8A-C). Ground squirrel cone ribbons were 824 ± 30 nm long (n = 29 ribbons; Fig. SD-H) and 225 ± 7 nm high (n = 25 ribbons; Fig. S8E,J) when fortuitously cut lengthwise and in cross-section, respectively. Based on a measured center-center spacing of 57.0 ± 0.9 nm (n = 116 vesicles Fig. S8J), we calculated that a two-sided rectangular ribbon held 114 SVs with 29 positioned at the base. We estimated the number of ribbon release sites in cone terminals from 3D STED images to be 22.1 ± 2.7 (n = 10 terminals from two retinas based on close associations between Ribeye, bassoon, and GluA4; data not shown), giving a total of 640 vesicles in the membrane associated pool. The close correspondence between the physiologically measured RRP of 600 vesicles and the ultrastructurally measured pool of 640 membrane-associated vesicles suggests that the short depolarizing pulses used to measure transmission at the cone synapse elicit release from the ribbon-docked pool.

We next used two approaches to estimate the maximal steady rate of vesicle replenishment at cone release sites. The first approach yielded a lower estimate by monitoring the change in membrane capacitance during a train of depolarizing pulses in the presence of high intracellular EGTA (10 mM). The short pulses repeatedly emptied the RRP while EGTA limited the domain of elevated Ca^2+^ to the base of the ribbons where Ca_v_ channels are located (Burrone et al., 2002; Zenisek et al., 2004; Beaumont et al., 2005; Van Hook and Thoreson, 2015). A linear fit to the average increase in C_m_ gave a replenishment rate of 17 vesicles-ribbon^-1^-s^-1^ (Fig 8C). A second approach used depolarizing steps of up to 1 s in length and established a rate of refilling under continual Ca^2+^ entry (Fig 8D; 37.6 ± 3.7 fF-s^-1^, n = 8 cones) equivalent to a turnover rate of 49 vesicles-ribbon^-1^-s^-1^. The results suggest that continuous Ca^2+^ entry accelerates turnover by 2.9-fold relative to that during intermittent Ca^2+^ entry (Babai et al., 2010). For modeling, we adopted a replenishment rate midway between the two numbers: 35 vesicles-ribbon^-1^-s^-1^.

We next used a Monte Carlo simulation to obtain a measure of the effective spread of glutamate in the synaptic cleft following vesicle fusion (Fig S8D,E). The simulation assumed that a vesicle contained 3000 glutamate molecules. At 200 nm from the release site, representative of the distance to receptors on a cb2 cell invaginating dendrite, glutamate concentration peaked at 510 µM. For comparison, at 600-800 nm from the release site, a range encompassing the locations of basal contacts midway between invaginations, concentrations peaked at 10-25 µM. Within 1 ms after fusion, the glutamate concentration decayed to around 10 µM or less at all sampling sites (Fig S8E). Since the EC_50_’s for Off bipolar cell AMPA and kainate receptors are typically >200 µM, receptor activation should be limited to the region around an invagination (i.*e.*, within 800 nm of the fusion site) and to brief (<1 ms) durations. Given an average rate of one vesicle fusion every 30 ms per ribbon (*i.e.*, ∼35 vesicles-ribbon^-1^-s^-1^), the Poisson distribution-governed probability that two events randomly occur at the same invagination within a 1 ms summation window, ∼0.0005, is equivalent to a rate of ∼0.5 s^-1^ or ∼10 s^-1^ over the entire terminal (∼20 invaginations). Using the trigonal synapse model (Fig 3F), a simple calculation suggests that the contents of two vesicles overlap an any contact at a rate of ∼60 events-s^-1^. Thus, under maximal steady release conditions (∼600 vesicles-s^-1^ per cone), fewer than 10% of independent event profiles overlap. The results and calculations suggest that probabilistic overlap of vesicular glutamate gradients is uncommon under maximal steady release conditions. However, this calculation ignores multiquantal release (*i.e*., simultaneous, highly localized fusion events involving the contents of multiple vesicles) which occurs at the ground squirrel cone synapse (Mehta *et al*., 2013).

### Light responses

Differences in the threshold responses of the Off bipolar cell types should be most evident when cones are hyperpolarized and vesicle fusion rates are low, a condition that occurs in steady bright light. We applied a bright light to suppress cone transmitter release and then decremented intensity to depolarize cones and enhance release. To reduce variability, we simultaneously recorded from nearby cb1 and cb2 cells. Using this strategy, we performed two types of experiments. In the first type of experiment, we saturated the cone response with 150 ms steps of bright light (Fig. 9A). In the dark prior to the light step, both the cb1a and cb2 cell displayed noisy inward currents due to continuous glutamate release from cones. A bright light stimulus hyperpolarizes presynaptic cones and stops release resulting in a dramatically reduced inward current and synaptic noise in both bipolar cells. After a saturating flash, cone membrane voltage slowly recovers to the resting state (Nikonov et al., 2006). During the slow return to baseline, plots of bipolar cell mean current and current variance, both measures of synaptic activity, (Fig 9B, upper and lower panels, respectively) showed a faster recovery in the cb2 than in the cb1a cell. Effective unitary response amplitudes (Fig 9C) were calculated during epochs 1-2 and 4-7 by measuring the current and variance change relative to epoch 3 and applying Campbell’s Theorem (see methods), which relates the ratio of the signal variance and mean to the size and shape of the unitary event (DeVries et al, 2006). We obtained canonical event shapes from small evoked synaptic responses. For the cb2 cell, the effective unit size at the start of the recovery (epoch 4) was reduced by 38.8 ± 11.1% relative to the size in the dark prior to the flash, transitioning to a 64.1 ± 17.8% increase relative to control in epoch 7. This increase is probably not related to receptor recovery from desensitization insofar as desensitization is minimal in cb2 cells during steady release and cb2 cell AMPA receptors recover from desensitization with a τ <20 ms (DeVries, 2000). In the cb1a cell, unitary responses were decreased by 82.8 ± 10.0% relative to the dark level during epoch 4 (significantly different from the decrease in the cb2 cell response, p < 0.0255; p < 0.0002 and p < 0.0007 for epochs 5 and 6, respectively, unpaired t-test), returning to the original level by epoch 7. The reduction in normalized effective event size was greater in the cb1a than in the cb2 cell during each phase of the recovery. Additionally, on the idea that cb1a cell kainate receptors are partially desensitized during steady release, a period of release suppression due to a bright flash should produce an increase in effective event size rather than the observed decrease.

**Figure 9.**
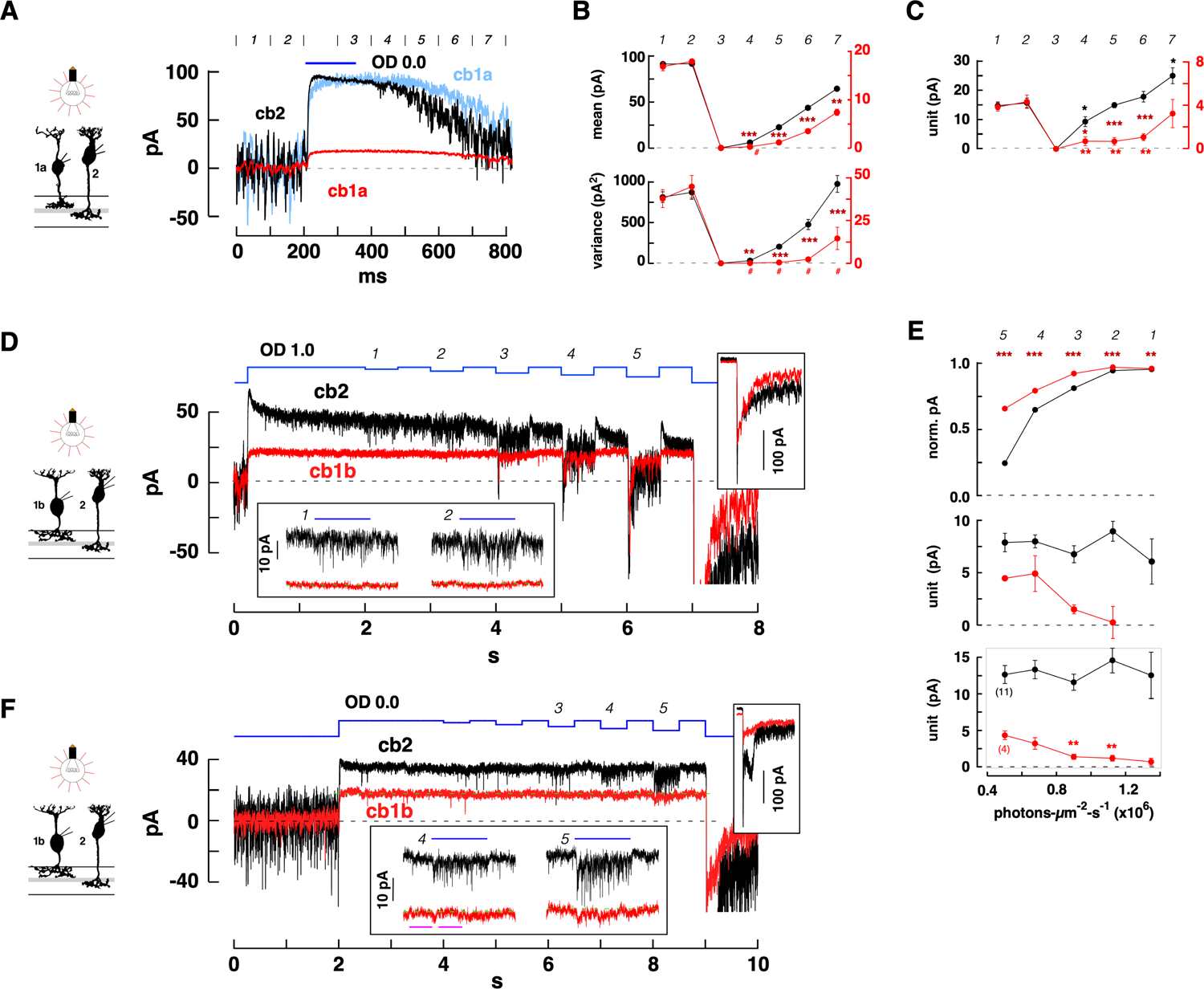
Threshold responses during light stimuli in Off bipolar cells. **A.** Responses in cb2 (black trace) and cb1a (red trace, original; light blue trace, scaled) cells simultaneously recorded during a 150 ms step of OD 0.0 light (1.575 x 10^7^ equivalent photons-µm^-2^-s^-1^). Epochs (duration = 100 ms) are marked by numbers (italics, above). Average of 4 trials. **B.** Average current (above) and variance (below; mean value ± S.E. obtained by separately analyzing each of the four responses; red and black symbols and axes correspond to cb1a and cb2 cell currents, respectively) for each epoch. Smaller S.E. bars are covered by plot symbols. cb1a and cb2 cell plots were normalized based on the average values in the first two epochs corresponding to the initial levels in darkness. Maroon (combination of black and red) asterisks indicate a significant difference between the normalized responses (0.05 > * > 0.01 > ** > 0.001 > *** > 0.0001; unpaired t test). The absence of an asterisk indicates a *n.s*. change. The red hashtags indicate that the cb1a responses were not significantly different from zero (one-sample t test). Points not marked by a hashtag are significantly different from zero. **C.** Plot of effective unit amplitude versus epoch. Red and black asterisks indicate significantly different from the average values in epochs 1 and 2 for the cb1a and cb2 cell, respectively. **D.** Responses of simultaneously recorded cb1b and cb2 cells during a long step of OD 1.0 light that was interrupted by a series of decrements (average of 3 traces; stimulus sequence is shown in blue). *Lower inset:* Magnified view of the light responses during the first and second decrements. *Right inset:* Responses at light-off shown on the same time scale but with a compressed current axis for completeness. **E.** Above: The normalized change in membrane current was plotted against the light intensity during the decrement (*i.e.*, the ‘step-to’ light intensity with the result that larger decrements are plotted toward the left on the x-axis; background light, OD 1.0, equaled 1.575 x 10^6^ equivalent photons-µm^-2^-s^-1^). Current was normalized by setting the value immediately before a decrement equal to 1 relative to the resting current in the dark. S.E. bars, calculated from the average traces, were covered by the plot point symbols. Normalized steady cb1b and cb2 responses were significantly different during all decrements (maroon asterisks). Middle: Effective unit amplitude was plotted against light ‘step-to’ intensity. Mean ± S.E. was calculated by separately analyzing each of the 3 trials. Below: Aggregate unitary responses for n = 11 cb2 and 4 cb1a/b bipolar cells (mean ± S.E.). Asterisks indicate that the values of the cb1 cell unit were significantly reduced during decrements 2 and 3 relative to decrement 5. **F.** Decrement responses from a different cb1b and cb2 cell pair during a long OD 0.0 background step (average of 2; repetitions were limited by photopigment bleaching at bright light intensities). The cb2 cell response displayed prominent epscs in the absence of events in the cb1b cell (decrements 3, and lower inset decrements 4 and 5). Magenta bars indicate the before and during regions used to measure the activity change in the cb1b cell in response to decrement 4 (200 ms intervals; p = .9924; not significantly different).

In the second type of experiment, to further exclude a role for receptor recovery from desensitization, we applied a steady bright-light stimulus to suppress release and, after a 2 s interval to allow cb1 and cb2 receptors to recover from desensitization, initiated a series of stimulus decrements to slightly increase transmitter release. The light responses of cb1 and cb2 cells were simultaneously recorded and compared. Each of the light decrements, including the two smallest (Fig. 9D, lower inset), produced a significant increase in inward membrane current and response variance in the cb2 cell. In the cb1b cell, the first decrement produced neither a significant change in membrane current (p = 0.2396 not different from zero by a t test) nor an increase in current variance (the measured variance change was negative), whereas the second decrement produced a small, 0.67 ± 0.12 pA, but significant (p < 0.0304) inward current without a significant increase in current variance (p = 0.7109; unpaired t test). Subsequent decrements produced significant current and variance changes in both cells. When cb1b and cb2 cell responses during a decrement were normalized with respect to the maximal steady decrement response to darkness and compared (Fig. 9E, top panel), the cb2 cell response amplitude was significantly larger for each decrement. Effective unitary responses were calculated by comparing the mean change in current amplitude and variance during a decrement (Fig. 9E, middle panel). Calculated cb2 cell unitary responses were roughly constant at ∼7-8 pA for all decrements. In contrast, the cb1b unit could not be measured during the first decrement (*i.e.*, the change in variance was assumed to be zero given a slight negative shift). The calculated unit size was 0.23 ± 2.66 pA during the second decrement (*i.e.*, not different from zero), and then increased to almost 5 pA over the last three decrements. Plots of unitary event amplitude versus light intensity decrement (Fig 9E, lower panel) were also obtained by averaging data from both paired and individual recordings performed at the same background intensity. In the aggregate data, the effective cb2 unit remained relatively constant in relation to decrement size whereas the cb1 cell unit was significantly reduced during smaller light decrements. An additional paired recording was performed in the presence of a maximal steady light background that fully blocked synaptic activity in both a cb2 and cb1b cell (Fig. 9F). In this experiment, we observed vigorous epsc activity in the cb2 cell during decrement 4 contemporaneous with a statistical absence of activity change in the cb1b cell (*i.e.*, comparing the before and during intervals marked by the magenta bars; unpaired t-test; there was an initial small response transient). The results are consistent with the idea that bright light reduces or eliminates event noise in cb1 cells under conditions were signaling persists in cb2 cells.

## Discussion

At the mammalian cone photoreceptor terminal, approximately 20 ribbon synapses release their vesicular contents into membrane invaginations that open onto a common, compact basal surface. The transmitter gradient at the basal surface is then sampled by the postsynaptic bipolar cell types. In mammals, but not in lower vertebrates, Off- or ionotropic-receptor expressing bipolar cells almost exclusively make basal contacts. To study the role of these specialized contacts in graded signaling, we developed a method for comparing the number of vesicles released by a cone to the size of the epsc response in a postsynaptic Off bipolar cell (Szmajda and Devries, 2011). We identified two sampling principles. In the first principle, the Off bipolar cell types have different thresholds for transmitter release and function to different extents as coincident event detectors. cb2 cells make invaginating contacts close to ribbon transmitter release sites, an exception to the general rule for Off bipolar cells, and respond at the single event level. cb3 and cb1 cell receptors are more distant from release sites. cb3 cells failed to respond to most single events while cb1 cells responded exclusively to coincident events or multiquantal release. Due to a tight association between depolarization and vesicle fusion (Fig 8B), coincident events are more likely to occur during light intensity decrements that produce an abrupt depolarization in a cone. Above the cb1 cell threshold, the responses of all Off bipolar cell types are linearly related, consistent with results in the mouse (Franke *et al*., 2017). The second principle is based on the idea that while the mean rate of release at each of a cone’s ribbons is governed by membrane voltage, the timing of release at each ribbon is stochastic and independent of release at neighboring ribbons. In this view, a bipolar cell can improve its estimate of the cone signal by sampling from multiple ribbons up to the limit of the entire terminal population. Our results suggest that some cb2 and cb3 cells sample from most or all of a cone’s ribbons via local dendritic contacts. Surprisingly, above threshold, cb1a cells, which make a few contacts at the center of a cone terminal, may achieve broad sampling by responding in a range where multiple release events create overlapping gradients. This tiered threshold organization of the cone to Off bipolar cell synapses may maximize the information transfer per amount of released glutamate (Savtchenko et al., 2013).

### Detection properties of EAAT5 at the cone terminal

The cone glutamate transporter anion conductance provides a way to measure transmitter release that is convenient, sensitive, and readily paired with recordings of the postsynaptic bipolar cell response. However, measuring transmitter release with the transporter has three potential drawbacks. The first drawback is that transporters might fail to report some release events. However, if failures to detect release were common, then evoked bipolar cell epscs would frequently occur in the absence of cone tpscs, a phenomenon that was almost never observed (Fig 2B and 9 cb2 pairs involving more than 100 trials each). Additional evidence supports the idea that cone transporters report all release events: the onset of the transporter current and small evoked epsc are equally rapid (Szmajda and Devries, 2011) suggesting that transporters are located close to release sites; the EC_50_ for glutamate at EAAT5 is suitably low (Gameiro *et al*., 2011); and, STED images show that EAAT5 clusters are widely distributed at the cone terminal base (Fig 1D). The second drawback is that false positive responses can interfere with tpsc measurements, especially at high spontaneous release rates. We excluded recordings with high spontaneous rates. Evoking release allowed us to further exclude tpsc responses that fell outside of a tight temporal window. Third, cone transporters display a saturating nonlinearity that can obscure vesicle counts in the upper ranges. In our model, linear detection hinges on the idea that transporter clusters are broadly distributed over the terminal surface while quantal release at an invagination impacts only a local transporter pool. In this scenario, saturation is related to the probability that an invagination releases two or more vesicles onto the same pool of transporters at the same time. A simple calculation shows that, on average, approximately 10% of the invaginations release two or more independent vesicles when the total amount released by a stimulus is 10-12 quanta (assuming 20 ribbon release sites per terminal) rising to 26% when 20 vesicles are released. We adopted 10-12 released quanta as the limit of the linear range for analyzing the transporter signal. This level is consistent with the measure of saturation provided in Fig S1C.

### Synaptic mechanisms of cb2, cb3, and cb1 cells

A model of cleft diffusion (Fig S8D,E) suggests that the effect of glutamate following single vesicle fusion is localized to the region around the invagination. The model is supported by recordings from cb2 cells which make invaginating cone contacts. For cb2 cells, the outcome following vesicle fusion is either a rapid >6 pA epsc or a response failure. A failure can occur if transmitter is released into an invagination that lacks a cb2 cell dendritic contact. In this view, linearity results from response summation among effectively isolated invaginating contacts when the probability of release at a ribbon is less than one, and from the near linear relationship between glutamate concentration and response at cb2 cell AMPA receptors when the average number of vesicles released per invagination is ∼2-5 (Grabner *et al*., 2016). A similar linear response relationship was found between salamander rods and horizontal cells, which also make contacts within invaginations (Hays et al., 2020).

Cb3 cells failed to detect transmitter >65% of the time when a cone released a single vesicle. If detection in cb3 cells follows a similar relationship to that in cb2 cells, the failure rate should gradually transition to 0% when 10 or more vesicles are released by a brief depolarization. Instead, the failure rate in cb3 cells approached 0% when 3-5 vesicles fused. The paradox of low sensitivity during single vesicle fusion and high sensitivity following the fusion of 3-5 vesicles can be resolved if cb3 cell dendrites, as a group, sample release from nearly all of a cone’s invaginations and each dendrite responds to the summed glutamate profiles of two or more events from neighboring invaginations or ∼35% of unitary events that are either larger or nearer. Complete sampling of ribbon-mediated release is supported by STED images that show an extensive network of GluK1 labeling beneath each cone terminal (Fig 3G-I). Cb3 cell receptors are capable of responding to glutamate near the concentrations found at basal membrane sites: Scaling up the results from responses to low concentrations of glutamate (*e.g.*, to 25 µM glutamate; Fig S7A,C) by two-fold (*i.e.*, to ∼50 pA) so that the peak receptor response in 18 mM glutamate and the maximal epsc at a cone to cb3 cell synapse match (∼500 pA; e.g., Fig 3B, *inset*) gives an epsc response size of ∼2.5 pA per contact when divided among 20 contacts. Response amplitudes near the quantal level can be reduced by glutamate binding to local transporters, a saturable effect that could account for the increase in effective unit size during larger epscs. While averaged cb3 cell responses (Fig S3F) suggest authentic failures, we could not determine whether cb3 cell epscs had a minimum amplitude or whether a “soft” threshold contained a continuum of amplitudes down to zero (Fig 3C; Fig 5D,E). At the lowest transmitter levels, stochastic single channel transitions may complicate event detection. The benefit of an event amplitude that scales with release rate may be to reduce the impact of quantal noise on signaling during bright light, a period when the vesicle release rate is low and where small decrements in light intensity produce small responses that are optimally encoded by small events. (Fig 9D, decrement *2* and Fig 9E).

In comparison to cb3 cells, cb1a cells rarely responded to depolarizations in the cone-linear range. Consequently, power-law fits to the scatterplot were not significantly better than linear fits over this range. Notably, a line fit with a close-to-zero slope could signify linear sampling of only 1-2 release sites by cb1a cells, which STED images show end in the central region beneath the cone. We tested this possibility and found that it could not account for the amplitude of the peak epsc response. Since the cb1a cell epsc amplitude starts to grow in the same range as the cone tpsc saturates, we compared the simultaneous responses in cb1a and cb2 cells postsynaptic to the same cone. A threshold nonlinearity in the cb1a cells is directly supported by a positive y-intercept in scatterplots of cb2 versus cb1a cell peak epsc response (Fig 5A-C). The offset may result from a combination of glutamate binding at basally located transporters (modeled in Fig 7F, *right*) and a cb1a cell receptor with a high EC_50_ for glutamate (Fig 6, Fig S7A,D). During moderate cone depolarizations, the epsc amplitudes of cb1a and cb2 cells are linearly related. We hypothesize that the linear relationship occurs under conditions where release saturates glutamate transporter binding sites and where both cb1a and cb2 cell receptors respond to incremental increases in glutamate concentration with Hill coefficients of ∼1. The results are consistent with the idea that cb1a cells filter-out the quantized noise of transmission under steady illumination, while preferentially signaling transmitter release that is synchronized by light decrements. A threshold non-linearity is found at some cone to Off bipolar cell synapses in the salamander, but the mechanism is unknown (Schreyer and Gollisch, 2021).

### Alternative mechanisms for producing distinct responses in the Off bipolar cell types

We considered the notion that differences in bipolar cell threshold might result from presynaptic mechanisms, namely systematic differences in activation thresholds among individual ribbons (Grassmeyer and Thoreson, 2017) followed by segregated contacts with Off bipolar cell types. First, under gentle stimulus conditions, vesicle fusion is organized around probabilistic Ca^2+^ nanodomains that are associated with portions of a ribbon and do not extend between ribbons to produce correlated release (Grassmeyer and Thoreson, 2017). Therefore, when two Off bipolar cells are simultaneously recorded during presynaptic cone depolarization, substantial within-trial correlations indicate that they sample from a common set of ribbons (Fig S6). Second, instead of varying the amount of transmitter released at a cone terminal by changing the strength of a depolarizing pulse, we varied the amount of released transmitter by repetitively applying a maximal 1 ms stimulus at various inter-pulse intervals within the range of the time constant for vesicle refilling at terminal docking sites (τ = 0.7 – 1.0 s (Grabner *et al*., 2016)). During a strong stimulus, release should occur at all ribbons irrespective of Ca^2+^ channel number or V_1/2_. For both cb2 and cb3 cells, the relationship between response amplitude and stimulus frequency was fitted by an exponential curve (Fig S10). A 16 Hz stimulus produced observable responses in both cell types. In contrast, a plot of cb1a cell response versus inter-pulse interval had a prominent sigmoidal shape such that responses at stimulus frequencies higher than 2-4 Hz were severely attenuated. This attenuation is likely unrelated to the rate of recovery from receptor desensitization insofar as a major time constant for recovery is 30-fold faster in cb1 than in cb3 cells (Fig S7I,J). The attenuation is consistent with a postsynaptic threshold. Thus, a presynaptic mechanism is unlikely to account for the observed bipolar cell threshold differences.

### Signaling and multiquantal release

While threshold filtering reduces the responses of both cb1 and cb3 cells to individual quanta, both cell types continue to respond to multiquantal events. Multiquantal events are thought to result from either the simultaneous fusion of vesicles that are adjacent at a ribbon (Singer *et al*., 2004) or from the compound fusion of ribbon-associated vesicles either prior to or during a membrane fusion event (Matthews and Sterling, 2008; Moser et al., 2020). The selective sensitivity near threshold of cb1/3 cells to multiquantal events raises the possibility of a multiplexed code at the cone synapse where single quantal and multiquantal events are preferentially signaled by different Off bipolar cell pathways. Multiquantal release can extend the dynamic range of bipolar cell synapses in zebrafish in response to strong stimuli (James et al., 2019). Additionally, the fraction of multiquantal events may be regulated by changes in synapse protein composition (Hays *et al*., 2020) which raises the possibility of modulation over the time course of hours or days. However, a mechanism for differentially regulating the proportion of multiquantal events on a moment-to-moment basis during moderate stimuli remains unclear.

### Processing at cone synapses in other mammals

While vertebrate retinas share a general plan, specific neuron types, even among mammals, display adaptations to ecological niche (Baden et al., 2020). Ground squirrel cone photoreceptors have the fastest light responses among mammals (Cao et al., 2014), and flicker ERG responses also suggest that post-receptoral processing is fast (Jacobs et al., 1980). Consistent with rapid signaling, AMPA receptor-expressing cb2 cells make invaginating contacts with cones. While the invaginating contacts of the Off cb2 cell may find no direct analog in other mammals, the situation is different for cb1a/b and cb3a/b cells. Results from primate and mouse retinas support the idea that Off bipolar cells use two different types of kainate receptors. In the primate, one set of Off bipolar cells, consisting of the DB2 and DB3b, has receptor responses and pharmacology similar to that of cb3a and cb3b cells (DeVries, 2020), and also expresses GluK1 receptors by immunohistochemistry (Haverkamp et al., 2001; Puthussery et al., 2014) and transcriptomics (Peng et al., 2019). The kainate receptors in the three other types of Off bipolar cells, the flat midget bipolar (FMB), DB1, and DB3a, have a different set of properties. In these three types, responses quickly recover from desensitization and GluK1 labeling is not evident by immunohistochemistry (Puthussery *et al*., 2014). Of the three types, the FMB and DB1 cell receptors most closely resemble those of ground squirrel cb1a and cb1b cells. A similar division in synaptic receptor properties is found between types3b and 4 in the mouse, which are similar to cb3a/b cells in the ground squirrel, and types 1 and 2, which are similar to the cb1a/b in the ground squirrel (Puller et al., 2013; Ichinose and Hellmer, 2016). Thus, the strategy of distributing signals to Off bipolar cell types with different thresholds may also apply in the primate and mouse.

### Comparison with other systems

Threshold coding has been proposed in other sensory systems. Cochlear inner hair cells transduce stimulus intensity over a six order of magnitude dynamic range into graded changes in membrane voltage that are communicated to postsynaptic afferents at ribbon synapses (Liberman et al., 2011). There are 5-30 ribbon synapses in an inner hair cell depending on tonotopic location in the cochlea and species (Liberman et al., 1990; Pangrsic et al., 2018; Moser *et al*., 2020). Each ribbon synapse is spatially isolated from its neighbors and communicates with a single action potential generating afferent (Meyer et al., 2009; Ohn et al., 2016). Single unit recordings suggest that the sensory afferents each encode only a portion of the dynamic range: afferents supplied by ribbons on the pillar side of the hair cell have a high spontaneous spike rate and low threshold for signaling, whereas the afferents that receive input from modiolar-side ribbons tend to have low spontaneous firing rate and high threshold (Liberman, 1982). Systematic differences in active zone structure and function implicate a presynaptic origin for the different coding properties (Liberman *et al*., 2011; Ohn *et al*., 2016; Ozcete and Moser, 2021). Pillar active zones, associated with low threshold afferents, tend to have Ca^2+^ channels that activate at more hyperpolarized voltages, while the Ca^2+^ channels at modiolar active zones, associated with high threshold afferents, tend to activate at more depolarized voltages (Ohn *et al*., 2016). While the main site of processing may differ, presynaptic for inner hair cells and postsynaptic for Off bipolar cells, both appear to use a ‘divide and conquer’ strategy to encode a range of sensory input.

## Conclusion

Heinz Wässle has likened the layout of the cone terminal to an integrated circuit in its compactness and ultrastructural complexity. By using the cone transporter to quantitate transmitter release, our results support a view in which the glutamate gradient under steady and slightly varying light conditions roils the terminal surface, varying dynamically in space and time. The different Off bipolar cell types, by virtue of contact location, contact number, receptor properties, and position relative to EAAT5 transporters, sample this dynamic gradient in different ways with different tradeoffs. The invaginating cb2 cell trades a reduced dynamic range for speed of signaling (Grabner *et al*., 2016). cb1a cells trade sensitivity to local changes in glutamate concentration for the ability to encode the higher, global changes in concentration that result from strong visual stimuli. cb1a cells also eschew a multiplicity of energetically demanding synaptic contacts for one or a few contacts localized to the parabolic center of the terminal. cb3 cells sample from almost the entire population of ribbons, providing a high-fidelity response to the cone signal over a broad range, but lack the sensitivity of cb2 cells to quantal release. Together, these outputs cooperate to encode the mammalian cone light-off signal.

## Acknowledgements

This work was supported NIH R01 EY012141 and an unrestricted grant to the Dept of Ophthalmology from Research to Prevent Blindness (S.H.D.) and JSPS KAKENHI grant #19H01140 (K.K. and D.F.).

## MATERIALS & METHODS

### Preparation and Electrophysiology

All procedures were performed at Northwestern University and approved by the Institutional Animal Care and Use Committee. The procedure for making ground squirrel (*Ictidomys tridecemlineatus*) retinal slices has been described (DeVries et al., 2006; DeVries and Schwartz, 1999). For experiments involving light responses, retinal slices were obtained under dim red illumination (Saszik and DeVries, 2012). During recordings, slices were visualized with a Zeiss Axioskop-2FS microscope using a 63x water immersion objective under infra-red illumination. Recordings were made with Axopatch 200B amplifiers (Molecular Devices) and signals were filtered at 5 kHz and digitized at a rate of 10 or 16.6 kHz with a HEKA ITC-18 A/D board (HEKA Elektronik) operated with custom software (Igor Pro 6.1; WaveMetrics). Patch pipettes were pulled from borosilicate glass capillary tubes to tip resistances of 8-12 MΩ.

The external solution consisted of (in mM): NaCl 115, KCl 3.1, MgSO_4_ 2.48, glucose 6, Na-succinate 1, Na-malate 1, Na-lactate 1, Na-pyruvate 1, CaCl_2_ 2, and NaHCO_3_ 25, 0.05% phenol red, and was equilibrated with 5% CO_2_/95% O_2_ to a pH of 7.4. Picrotoxin (50 µM; Sigma, P1675) and strychnine (10 µM; Sigma, S0532) were included in the bath. The patch pipette solution for the bipolar cell contained (in mM): KCl 120, Cs-EGTA 10, MgSO_4_ 2, HEPES 10, ATP 5 and GTP 0.5; pH 7.35 with KOH. The cone pipette solution contained (in mM): KSCN 115, K-EGTA 10, MgSO_4_ 2, HEPES 20, ATP 5 and GTP 0.5; pH 7.35 with KOH. Intracellular solutions contained combinations of a non-fixable tracer, either Sulforhodamine 101 (Molecular Probes, S359) or BODIPY 492/515 (Molecular Probes, D3238) and a fixable tracer, either cascade blue hydrazide, trilithium salt (0.1 mM; Invitrogen, C3239) or 0.5% Neurobiotin Tracer (Vector Laboratories, SP-1120). Labeled cells could be visualized using immunohistochemistry followed by confocal imaging (Light et al., 2012) or in situ immediately after the recording with a Prime95B (Photometrics) camera controlled by µManager (micro-manager.org). All solutions were corrected to an osmolarity of 285 ± 5. Command voltages were not adjusted for liquid junction potential. Experiments were performed at 32 − 33 °C. Pharmacological agents include: UBP310 (Tocris, #3621), GYKI53655 HCl (Tocris, #2555), DL-TBOA (Tocris, #1223), and glutamic acid (Sigma, G1251). Membrane capacitance was measured with a HEKA EPC-10 using the ‘sine+dc’ Lockin routine in Patchmaster software. The intracellular solution and recording details are the same as (Grabner *et al*., 2016). Retinas were stimulated by a light-emitting diode (574 nm) attached to a microscope video port. LED intensity was controlled by pulse-width modulation. Light sources were calibrated with a photodiode detector (International Light) that was positioned beneath the microscope objective. Light intensity was converted to photons at 520 nm, the λ_max_ for the ground squirrel green cone pigment (Kraft, 1988). Figure legends show light intensities. During light stimuli, unitary responses were analyzed using Campbell’s theorem (Rice, 1944),

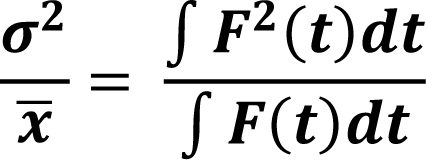

where ^-^*x* and σ^2^ are the signal variance and mean and the integrals are equal to event area and event area squared. Canonical events were obtained from cb1 and cb2 cells during 1 ms cone depolarizations. The event amplitude (*i.e.*, in the denominator) was scaled to find the unique value that matched the variance to mean ratio. Application of Campbell’s theorem assumes both linear event summation and that stochastically occurring events are the predominant source of noise in the difference trace. Both conditions may be approximate during bipolar cell light responses over a range of intensities.

### Acquisition and analysis of tpscs and epscs

Epsc and tpsc traces were Gaussian filtered in software with cutoffs of 1000-1250 Hz and 500-750 Hz, respectively, before further processing. Cones were stepped in voltage clamp from a holding potential of −70 mV to a level between −50 and 0 mV for 1 ms. Tpsc responses during an epoch (a train of steps to the same voltage) were measured as the average current during a 5 ms window that encompassed the inward current maximum relative to the average value during an 8 ms baseline prior to the step. Peak responses from one or more consecutive epochs were accumulated into amplitude histograms. Amplitude histograms were fitted by a sum of Gaussians with variables for individual peak amplitudes (including failures), mean ‘quantal’ size (Δx), and failure and success peak Gaussian widths, σ_f_ and σ_s_, respectively. The peak location of the failure distribution, x_0_, was unconstrained. The Gaussian widths for the failure and subsequent success peaks followed the formula (σ_f_^2^ + *n*(σ_s_^2^))^1/2^ where *n* is peak number. Each tpsc response from an entire run consisting of multiple epochs was assigned an effective quantal content, *m*, by binning according to x_0_ + Δx(*m* − 1/2), x_0_ + Δx(*m* + 1/2). Tpscs with *m* > 12 were assumed to have non-linear summation. All assignments were inspected for the presence of spontaneous cone transporter events either before or immediately after the 1 ms stimulus. In the case of overlap, amplitudes were either assigned manually or, in rare cases where assignment remained uncertain, the trial was discarded.

Epsc peak amplitude was determined for all Off bipolar cell types using the automated routines described in (DeVries *et al*., 2006). In brief, copies of traces were filtered in software (Gaussian filter with a cutoff frequency of 200 Hz). The baseline noise was calculated from time zero to the start of the 1 ms step for each filtered trace and the values for all traces were averaged to obtain σ_n_. The response threshold was empirically set to 3.5σ_n_, which was typically 0.8 − 1.5 pA. The unfiltered epsc responses in the series were then averaged and fitted over the response interval by least squares minimization with a function that could approximate both fast and large cb2 and slow and small cb1/3 events. The shape of the function, ConvExpDiffusion, was determined by convolving a fast exponential rise (τ = 0.1 ms) and decay (τ = 1.0 ms) with a profile obtained from an equation for diffusion from a point source in three dimensions. The variables of the diffusion equation, peak amplitude, diffusion radius, and start time, were saved. Each 200 Hz filtered trace was sorted into one of two groups, failures or successes, based on whether the epsc exceeded threshold during the response interval. For epscs that exceeded threshold, peak amplitude, start time, and rise time were determined for the unfiltered trace by fitting with ConvExpDiffusion. For responses that failed to exceed threshold, traces were fitted within the response interval using ConvExpDiffusion to determine positive or negative amplitude while using the saved values for radius and start time obtained from the series average. The fitting routine issued error messages for instances of poor fit, which could then be inspected for interference by spontaneous events or noise. The fitting routine was robust to small, but suprathreshold, cb1 and cb3 cell epscs whose rise and peak regions were interrupted by apparent single channel transitions.

For plots of percent failure versus cone quantal content, the confidence interval was based on a maximum likelihood estimation for coin tosses. For a biased coin with an intrinsic probability of heads, *p*, the binomial distribution gives the probability of observing a specific number of heads, *h*, in *n* trials. The cone to Off bipolar cell synapse presents the inverse problem where *h* (i.e., the number of failures) and *n* are known, and the task is to estimate the likelihood that the intrinsic probability, *p*, is within a certain range of values. The relative likelihood of an intrinsic probability *p* given *n* and *h* is obtained from the beta probability distribution with parameters *h* + 1 and *n* – *h* + 1 such that the greatest likelihood occurs at *p* = *h*/*n* and the confidence interval is bounded by the probability values that exclude the lower and upper 25% of the area under the curve, respectively (*i.e.*, the 50% confidence interval). Sample sizes of 2 or smaller were omitted from the plot due to the large uncertainty associated with the measurement.

### Rapid perfusion of somatic patches

A rapid perfusion pipette was mounted on piezoelectric actuator (Burleigh, PZS-200) and driven by an amplifier (Burleigh, PZ-150 M). The command voltage for the amplifier step, typically 60 ms in duration, was filtered in software by convolution with a Gaussian (width = 1.2 ms) to damp oscillations. Four-barreled glass (Vitrocom, Mountain Lakes, NJ) was mounted on a Kopf vertical puller and pulled so that each square barrel had a width of ∼100 µm. The two side barrels were sealed with Sylgard. Solutions were fed under pressure to the two central barrels via small bore polyimide-coated quartz tubes (Polymicro Technologies, Phoenix, AZ). The flow into one barrel contained control solution and had a single input. The flow into the other barrel could be switched among five test solutions. Control solution contained (in mM): NaCl 125, KCl 3.1, MgSO_4_ 1.24, CaCl_2_ 2, and Na-HEPES 10. pH was adjusted to 7.4 with NaOH and osmolarity was adjusted to 285 ± 5 mosm with NaCl. Na-glutamate (18 mM) was substituted for NaCl to make the stock test solution. Serial glutamate dilutions were made from the stock. Junction potential differences between the control and 18 mM test solutions were used to measure open tip switching time at the end of an experiment.

To withdraw bipolar cell somata, all adherent processes from other bipolar cells were first removed from an isolated bipolar cell soma via a suction-clearner pipette. Whole cell access was obtained with a pipette filled with bipolar cell intracellular solution containing sulforhodamine 101 and Neurobiotin Tracer. Continuous negative pressure was applied to the pipette by syringe and the soma was slowly withdrawn from the slice leaving behind a soma-less but otherwise intact cell remnant which could be identified under epifluorescence or, after fixation, by confocal microscopy.

For glutamate uncaging, 5 mM MNI-caged-L-glutamate (Tocris, #1490) was added to the extracellular solution and applied to the synapse by a local puffer pipette. Bipolar cells did not respond to puffer-applied caged-glutamate in the absence of the flash. Spots, ∼3 µm in diameter at the level of the slice, were delivered by a Vortran Stradus Laser (405 nm, nominal power = 50 mW) via a Rapp OptoElectric Spot Illumination System that was attached to the epifluorescence port. Laser power was set using Vortran Stradus Control Software and an electronic shutter was driven by a TTL pulse from the ITC-18. The microscope objective was an Olympus LUMPlanFLN 60x/1.00W.

### Modeling vesicle fusion at the cone synapse

#### Cb2 cell model

The model cone terminal contained 20 invaginating transmitter release sites of which *x* were contacted by dendrites. The probability of response failure, *p_f_*, was calculated as a function of the number of simultaneously released cone quanta, *n*, for each *x* between 1 and 20. Every quantum had an equal probability of release at all 20 sites. For the case where a success occurs whenever a single vesicle is released at a site that is occupied by a dendrite, the probability of failure *p_f_*(*n*) = (1 − *x*/20)*^n^*. Probability curves were calculated for each *x* and associated with a cb2 cell data set by least squares minimization. *p_f_*(n) plots at a given *x* consisting of discrete points were fitted with an exponential decay curve for display (Fig 2F).

#### Cb1/3 cell model

The model cone terminal had 19 invaginating release sites arranged in a trigonal array (Fig 3F). Contact number could be varied from 1 to 24 with individual contacts located at the centers of triangles whose vertices are release sites. A response rule was constructed and, for a given number of dendritic contacts, the relationship between the failure probability *p_f_* and the number of released cone quanta, *n*, was determined by running a simulation. In a typical response rule, a success occurs when any one dendritic contact receives 2 or more (or ≥3, ≥4, or ≥5) quanta in any combination from the 3 surrounding vertices. Computationally, in a trial, each bin in an array of 19 release sites is populated with *m* release events where *m* = 0, 1, 2, 3 … as determined by the Poisson distribution and a random number generator. The Poisson average in a simulation was set to *n*/19 where *n* is the user assigned total number of quanta released in the trial. To determine the *p_f_* for *n* cone quanta, the program generated 1000 trials while only retaining those where the sum of all released quanta across bins equaled *n*. For example, to determine *p_f_* for the condition in which a cone terminal releases exactly 2 quanta, the sum across all retained bins must equal 2. The retained ‘release site’ trial array was then used to populate a 24-bin ‘dendrite’ trial array using the mapping from Fig 4F and the response rule. For example, for the ≥2 rule above, dendrite array bin 1 would be set to success (= 1) if the sum of release site array bins 2, 3, and 6 is ≥2 or failure (= 0) otherwise. An analogous operation is then performed for each of the 24 dendrite bins. For each retained trial, if the sum of the dendrite array is 0, then the trial is a failure, otherwise it is a success. The proportion of failed trials was then calculated to give *p_f_* for the number of cone quanta released. The simulation was repeated for *n* = 1… 20 to obtain the entire probability curve. The role of contact number was examined by masking the dendrite array so that the values in the first *x* bins were maintained while values in the remaining bins were set to 0. To model a certain percentage of success at the single quantal release level while otherwise maintaining the ≥2 response rule, 20-35% of the bins that contained 1 vesicle in the retained array were randomly tagged so as to guarantee a success when mapped to a bin in the dendrite array. To simulate multiquantal release, each of the *m* events in each release array bin was associated with a Ca^2+^ channel opening that could randomly lead to mono-vesicular release 80% of the time and di-vesicular release 20% of the time. The total release events in the bin were adjusted accordingly. For trial retention, the sum of all quanta still had to equal *n*. Hence, multiquantal release does not contribute to *p_f_* when the model cone releases only a single quantum (*n* = 1). Fits were evaluated by least squares minimization against data points and the best fit was arrived at through an iterative procedure.

### Processing tissue for IHC and STED microscopy

Tissue was either fixed in 4% PFA using standard procedures (Light *et al*., 2012) or in 2% glyoxal solution (Richter et al., 2018). Glyoxal fixative (10 ml) contained 7.325 ml distilled H_2_O, 2 ml absolute ethanol, 0.5 ml glyoxal (40% in water, Sigma-Aldrich, #128456), 0.075 ml glacial acetic acid, and 0.1 ml Na-acetate (3M, Ambion, #AM9740) to give a pH of ∼4.0. Pieces of freshly dissected retina with pigment epithelium, 1 x 1 mm, were placed ganglion cell side down onto dry Millipore filter paper (8.0 µm MCE Membrane, #SCWP02500) in a small petri dish. Cold PBS was added, and the pigment epithelium was peeled from the paper-adherent retina. The petri dish was placed on ice for 5 min and then cold glyoxal solution was substituted for the PBS. After 2 hrs on ice, the tissue was placed in the refrigerator at 4°C for 2 days. Tissue was removed from the refrigerator was washed 3x for 20 min each with cold DPBS (Gibco, #21600-010), and sliced with a tissue chopper into 100 µm or 300 µm thick sections for cross-sectional or whole mount views, respectively. Individual slices were transferred to MatTek 3 mm glass bottom culture dishes and incubated for 1 day in block solution containing 0.5% Triton X-100, 0.1% Na-azide, and 3% donkey serum 0.1 mM phosphate buffer. Whole mounts were treated with primary antibodies for 6 days (3 days for slices) in block at 10°C with gentle shaking. The tissue was then washed 6x for 30 min each and incubated in secondary antibody solution with block for 4 days (2 days for slices) with shaking at 10°C. The tissue was then washed 6x for 30 min each with 0.1 M phosphate buffer plus 0.1% Na-azide in preparation for mounting. The tissue was mounted in ∼1 g melamine resin consisting of 0.6 g 2,4,6-Tris[bis(methoxymethyl)amino]-1,3,5-triazine (melamine, TGI America, #T2059), 80 mg citric acid monohydrate (Sigma-Aldrich, #C1909), 20 mg of 8,000 MW polyethylene glycol (2%, w/w, Sigma-Aldrich, #89510), 5 mg caffeic acid (Sigma-Aldrich, C0625), 5 mg propyl gallate (Sigma-Aldrich, #02370), and 0.3 ml distilled water. The mixture was liquified by vortexing and then incubating on a horizontal shaker (200 rpm) in an oven at 55°C for 1 hr. For mounting, the tissue was rapidly washed 2x with distilled water that was then completely removed. The tissue was bathed in melamine resin for 1 hr at room temperature and then removed and transferred with a fine spatula to a glass slide. 15 µl of free resin was added to the tissue followed by coverslipping (Zeiss, High Performance, 18 x 18 mm, #1½). The resin was cured at 55°C for 2 days after which it had a refractive index of 1.52.

Primary antibodies, sources, and dilutions are listed below. Secondary antibodies (JacksonImmuno) raised in donkey against mouse (#715-005-151), rabbit (#711-005-152), goat (#703-005-155), guinea pig (#706-005-148), and chicken (#705-005-147) were reacted with NHS-ester dyes using standard procedures (http://abberior-instruments.com/wp-content/uploads/0236_20120316-labeling_protocol.pdf). NHS-ester dyes were ATTO 532 (Atto-tec, #AD 532-31), abberior STAR 580 (Abberior GmbH, ST580-0002), abberior STAR 635P (ST635P-0002), and CF680R (Biotium). Degree of labeling and final antibody concentration was determined with a spectrophotometer using published values for maximal absorption wavelengths, extinction coefficients, and correction factors (280 nm). Secondary antibody dilution was 1:100.

Anti-PSD95 PDZ domain, SySy 124014, guinea pig Anti-SLC1A7 (EAAT5), Sigma, HPA049124, rabbit Anti-CtBP2 (C-16), Santa Cruz Biotechnology, sc-5967, goat (1:100) Anti-GluR4, EMD Millipore, AB1508, rabbit (1:200) Anti-GluR5 (E-12), Santa Cruz Biotechnology, sc-393420, mouse monoclonal (1:500) Anti-ChAT, Chemicon, #AB144P, goat (1:100) Anti-GFP, Abcam, ab13970, chicken, (1:1000) Anti-Bassoon, SySy 141016, chicken, (1:500) Anti-Alexa Fluor 405/Cascade Blue, A-5760, rabbit (1:500)

### STED microscopy image acquisition and analysis

Super resolution microscopy was performed on a Leica SP8 3D-STED system using a 100X NA 1.4 oil objective, white light laser for excitation, 775nm depletion laser, and z vortex set for isotropic voxel generation under control of LASX software. 3D capture used 20 nm XY and 50nm Z steps using 16kHz scan rate, 8 or 16 line average and varied frame accumulation to build up signal. Image data were denoised using FIJI CSBDeep noise2void trained on each day’s data (https://imagej.net/plugins/n2v). Denoised images were deconvolved in Imaris 9.8.0 using the ClearView module. Deconvolution parameters were: Robust (iterative), 2.0 pre-sharpening gain, 10 iterations, denoising filter equal to 0.7. Specimen and medium refractive index were both 1.52. Unless otherwise noted, images are maximum intensity projections.

The analysis procedure in Fig 3G-I and Fig S4 had four steps. In step 1, square regions centered around each terminal’s receptor labeling were excised from a binarized (Fiji, default settings) STED image. The size of the square region was 4 μm x 4 μm. Excised images were superimposed and averaged to obtain the spatial profile of GluK1 or labeled bipolar cell dendrites beneath the terminal. Averages were converted to 8 bit. Step 2 involved feature extraction by Fourier filtering. A two-dimensional Fourier transform was applied to the average profile, the frequency components with low power were removed, and then an inverse transform was performed. The power cutoff threshold was determined manually for each image and is indicated in the figure panels. In step 3, contour borders were abstracted from the filtered profiles using a Canny edge detection algorithm and a Hough transform, the latter by assuming a circular or elliptical profile. Canny edge detection and Hough transforms were executed using Python OpenCV modules. The fourth step was to quantify the spatial bias of GluK1 labeling or dendrite distribution using moment analysis. Within the region formed by central and peripheral contours, the centroid coordinates were calculated twice, first for a uniform distribution and then for the actual intensity distribution. In both cases, the centroid coordinates (*Cx*, *Cy*) were given by the following equation:

where *xi*, *yi* are the *i*-th pixel coordinates of the formed region, and *R*(*x*, *y*) is the pixel value (*e.g.*, the overlap fraction of GluK1). The difference vector between the coordinates obtained from the uniform profile and actual data gives the magnitude and orientation of the bias of the GluK1 or dendritic process distribution.

### Receptor modeling

The glutamate receptor models for cb1a and cb2 were based on the Markov kinetic scheme proposed by (Hausser and Roth, 1997). The behavior of the receptors was represented by transitions between nine discrete states (Fig. S8A). The transition rates were different depending on the cone bipolar cell type (Table S3). The rates used for the cb2 receptor model were from (Grabner *et al*., 2016). The rate constants for the cb1a cell receptor model are new and were obtained using Particle Swarm Optimization (PSO) (Kennedy and Eberhart, 1995). The model output was optimized against receptor responses and parameters during various test glutamate applications (*i.e.*, response rise and decay, EC_50_, IC_50_, recovery τ’s from desensitization, and the percent open probability during long exposures to glutamate; Fig 6A-D, Fig. S7, Fig S8A-C) using a weighted sum of the absolute errors of the real and model data. The optimization process searched for the rate parameter set that minimized the weighting function. PSO was performed by using PySwarms. Maximal channel open probability was estimated using non-stationary fluctuation analysis from repeated responses of cb1a receptors to an 18 mM glutamate step administered by rapid perfusion. Timing jitter between separate responses in the 20-40 µs range was removed before processing by aligning the traces by their rising phase (data not shown).

### MCell synaptic modeling

We use MCell 3.4 and CellBlender 1.0.1 to construct a model synaptic cleft (Fig S8D) (Stiles et al., 1996; Stiles and Bartol, 2001). The synaptic cleft consisted of 20 μm x 20 μm planes corresponding to the pre- and post-synaptic membranes. The planes were separated by 16 nm (Lasansky, 1969; Raviola and Gilula, 1975). Glutamate molecules were released from a point in the center of the presynaptic plane with a diffusion coefficient of 400 μm^2^/s (Grabner *et al*., 2016). There is substantial uncertainty about the number of glutamate molecules in a vesicle in the range of 3000 – 8000 (Wang et al., 2019). Therefore, instead of releasing glutamate molecules in increments of “quanta”, we varied transmitter release in increments of 1000 molecules. The presynaptic plane was impermeable to glutamate while the postsynaptic plane was 10% permeable so that lateral glutamate diffusion in the cleft is consistent with previous work (Grabner *et al*., 2016). Permeability across the postsynaptic plane is consistent with it being composed of dozens of individual dendritic contacts each separated by an extracellular space. The planar distances of transporters and receptors from the release site was increased to simulate the effect of transmitter flow into the plane through an invagination-like cylinder 200 nm deep by 50 nm wide. Adding a cylinder instead made little difference in the outcome of the simulation. An annular patch (0.02 μm^2^) with 100 cb2 receptors surrounded the release site at a radius of 100 nm, and a square patch (0.04 μm^2^) with 350 Cb1a receptors was centered 600 nm away from the release site on ‘basal’ membrane. In addition, a fan-shaped patch with 3200 EAAT5 transporters was placed between those two receptor patches in a region of the model surface corresponding to basal membrane. The transporter model was based on a simple transition between glutamate bound and unbound states in the range of the measured EC_50_ for EAAT5. The forward and backward rates were 10^7^ M^-1^s^-1^ and 200 s^-1^, respectively. In some simulations, the transporter patch was moved to the side of the invagination opposite to the receptor patch and in others it was moved in the same direction but beyond the receptor patch.

---

**Table S1.** Spatial profile of GluK1 distribution beneath cone terminal. The column items are the features of the elliptical profile obtained from the five STED images shown in Figure S3B. “Circularity” is the ratio of the minor axis to the major axis of the detected elliptical profile; the closer the ratio is to 1, the closer it is to a perfect circle. The intensity in a pixel is given by the fraction of the summed images that contain a process at that position.

| Data label | Circularity | Pixel average (%) in |  |
| --- | --- | --- | --- |
|  |  | annulus | dimple |
| #0 | 0.767 | 60.92 | 49.79 |
| #1 | 0.871 | 67.50 | 53.74 |
| #2 | 0.821 | 85.58 | 83.40 |
| #3 | 0.982 | 73.69 | 67.13 |
| #4 | 0.963 | 67.74 | 61.00 |
| Mean | 0.881 | 71.09 | 63.01 |
| SD | 0.092 | 9.28 | 13.21 |

**Table S2.**
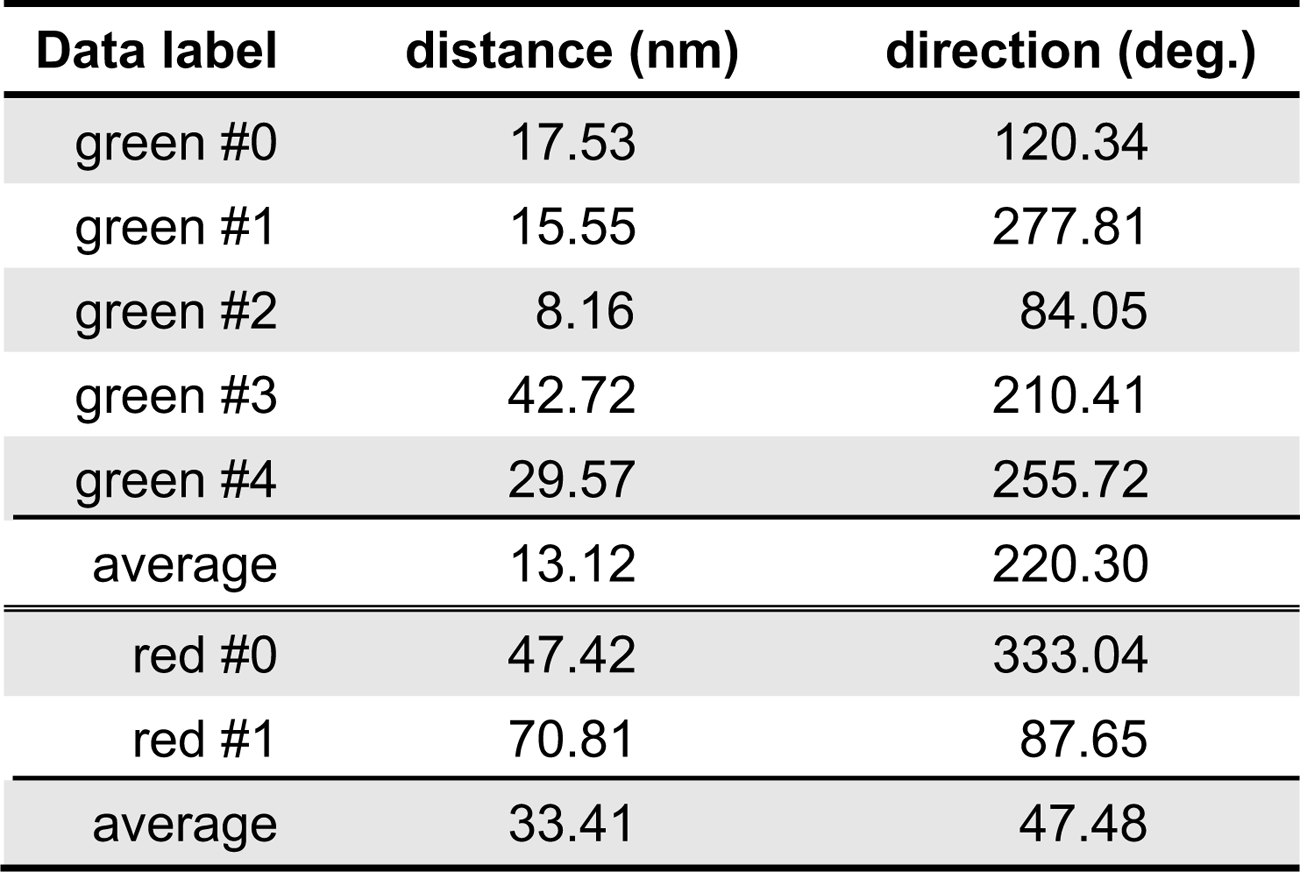
Centroid displacement. The data labeled “green” and “red” correspond to the filled-in green and red circles in Fig. 3I. The data labeled “average” corresponds to the open circles in Fig. 3I.

| Data label | distance (nm) | direction (deg.) |
| --- | --- | --- |
| green #0 | 17.53 | 120.34 |
| green #1 | 15.55 | 277.81 |
| green #2 | 8.16 | 84.05 |
| green #3 | 42.72 | 210.41 |
| green #4 | 29.57 | 255.72 |
| average | 13.12 | 220.30 |
| red #0 | 47.42 | 333.04 |
| red #1 | 70.81 | 87.65 |
| average | 33.41 | 47.48 |

**Table S3.**
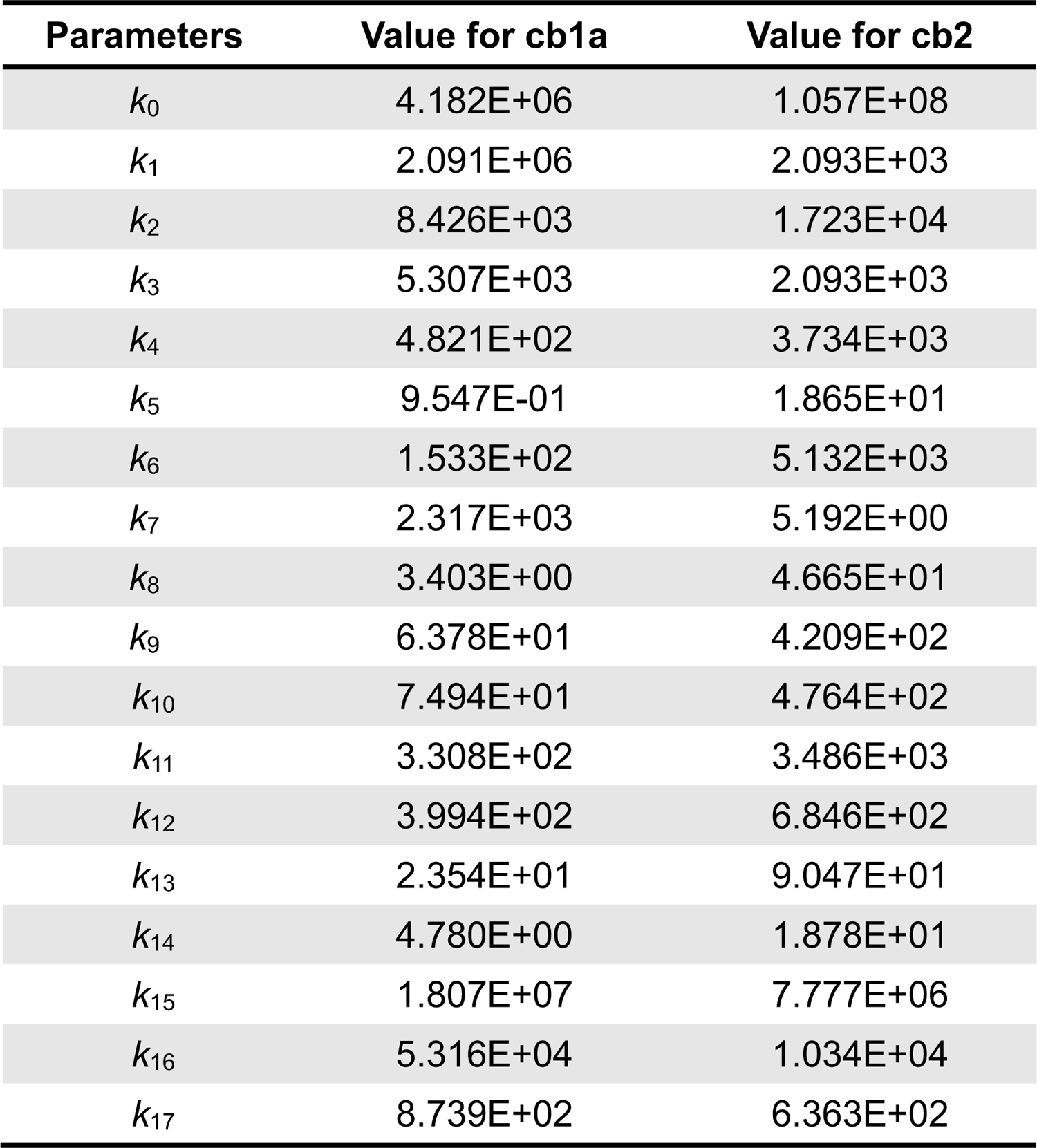
Rate constants in cb1a and cb2 models. The unit of the rate constants is M^-1^s^-1^ or s^-1^. In order to establish the microscopic reversibility of the model, let *K*_1_ = (*k*_7_\**k*_1_\**k*_6_\**k*_11_)/(*k*_15_\**k*_8_\**k*_3_), *K*_2_ = (*k*_5_\**k*_9_\**k*_8_\**k*_2_)/(*k*_4_\**k*_6_\**k*_10_) and *K*_3_ = (*k*_14_\**k*_17_\**K*_2_\**k*_12_)/(*k*_13_\**k*_5_\**k*_16_).

| Parameters | Value for cb1a | Value for cb2 |
| --- | --- | --- |
| $k_0$ | 4.182E+06 | 1.057E+08 |
| $k_1$ | 2.091E+06 | 2.093E+03 |
| $k_2$ | 8.426E+03 | 1.723E+04 |
| $k_3$ | 5.307E+03 | 2.093E+03 |
| $k_4$ | 4.821E+02 | 3.734E+03 |
| $k_5$ | 9.547E-01 | 1.865E+01 |
| $k_6$ | 1.533E+02 | 5.132E+03 |
| $k_7$ | 2.317E+03 | 5.192E+00 |
| $k_8$ | 3.403E+00 | 4.665E+01 |
| $k_9$ | 6.378E+01 | 4.209E+02 |
| $k_{10}$ | 7.494E+01 | 4.764E+02 |
| $k_{11}$ | 3.308E+02 | 3.486E+03 |
| $k_{12}$ | 3.994E+02 | 6.846E+02 |
| $k_{13}$ | 2.354E+01 | 9.047E+01 |
| $k_{14}$ | 4.780E+00 | 1.878E+01 |
| $k_{15}$ | 1.807E+07 | 7.777E+06 |
| $k_{16}$ | 5.316E+04 | 1.034E+04 |
| $k_{17}$ | 8.739E+02 | 6.363E+02 |

## Notes

### Competing Interest Statement

The authors have declared no competing interest.

