## Supplemental Figures for "Design of the mammalian cone photoreceptor to Off bipolar cell synapse"

**Figure S1:** Properties of the cone tpsc response. **A.** Histogram of the maximal cone tpsc responses to a 1 ms depolarization (left) in the set of 55 cones and the unitary response amplitudes (right) obtained from a subset of cones (black open bars) by fitting their amplitude histograms (15.5 ± 5.0 pA, mean ± S.D.; n = 24). The sample of 24 cones was obtained after the following exclusions: In the set of 55 cones (red bars; peak epsc = 409.0 ± 137.1 pA), 24 (black open bars; 420.1 ± 113.3 pA) were further analyzed. Reasons for exclusion from the amplitude histogram analysis include: 1) no response failures in the sequence (13 cones); 2) a high spontaneous event rate that caused evoked and spontaneous events to frequently overlap (10 cones); and 3) low maximal peak amplitude (2 cones; <200 pA). For six additional cones, the amplitude histogram fits were indeterminate. **B.** Category plots comparing the estimates of unitary event amplitude obtained by the different methods: amplitude histogram, $P_0$, and variance to mean (v/m) ratio. Colored lines connect results from the same cone. Letters denote the results from analyzing the trains in Fig 1F, H, and *inset* (above right). Points were plotted without (left) and with (right) normalization to the values obtained from the amplitude histogram. The variance to mean ratio occasionally reported larger than expected unitary event amplitudes because of infrequent but large multiquantal events. **C.** Plot of tpsc variance to mean ratio versus normalized tpsc amplitude for 13 cones (data from each cone is denoted by a different symbol; normalization was against the peak tpsc amplitude for the cone; the peak tpsc for the 13 cones was 465.0 ± 29.0 pA; mean ± S.E.). A sigmoid fit to the aggregate data had a maximum value of 22.7 pA and was constrained to approach zero at high mean values. The sigmoid fit had a ½ decay point at 54% of the mean response and a decay exponent of 4.9. The curve crossed the 90% level (red dashed line) at 34.2% of the maximal response. This percentage corresponds to a current of ~160 pA relative to the mean peak tpsc value of 465 pA, which is equivalent to ~10 average (*i.e.*, 15.5 pA) quantal units. Arrows denote the points (upright filled triangles) associated with the sets of traces that show the responses of a single cone during minimal (*inset*, above) and saturating (*inset*, below) 1 ms pulses. The response shown in red is a spontaneous tpsc that immediately followed a failure. The 13 cones in the plot were selected from the 24 in the sample based on having sufficient data points at intermediate mean current levels to provide information on the sigmoid decay.

**Figure S2: A.-E.** Scatterplots for 5 additional cone to cb2 cell pairs. Data points are averages over 10 pA bins of cone epsc amplitude. The standard deviation for the bipolar cell epsc peak is shown. Points that lack error bars result from a single measurement. Point location on the x-axis is also an average value relative to the 10 pA bin width. Least squares fits to linear (black) and power-law (cyan) equations are shown over the cone-linear range. Sampling percentage is calculated from the linear slope after normalizing for cone and cb2 bipolar cell unit amplitude. Circle colors correspond to those in Fig 2F,G and Fig S2F. **A.** Linear fit preferred, p = 0.3126 (F test). **B.** Power-law fit had an exponent of 0.69 and was omitted. **C.** Power-law fit preferred, p = 0.0004, exponent = 1.58. **D.** Slight preference for the power-law fit, p = 0.0123, exponent = 1.41. **E.** Linear fit preferred, p = 0.1453. Insets show the maximal cone tpsc and cb2 cell epsc for each pair. **F.** Percent failures versus cone quanta plots and fits for three of the pairs.

**Figure S3:** Properties of the cb3 cell responses during cone depolarization. **A.** cb3 epscs have the same time course irrespective of size. Epsc responses from a single pair (n = 3 total pairs with similar results). (left). Traces from several stimulus epochs were sorted into amplitude bins and averaged. (right). Normalized responses have nearly identical time courses. This scenario addresses the possibility that large epscs are large because they contain events that occur close to dendritic contacts, and hence are faster than smaller more distant events which contribute to small epscs. Same pair as Fig 3A,B. **B.-E.** Scatterplots for 4 additional cone to cb3 cell pairs. Power law fits (black curves), exponents ('b'), and corresponding data points in Fig 3E are indicated. Insets show the maximal cone tpsc and bipolar cell epsc. **F.** Analysis of cb3 cell response categorized as failures for evidence of small epsc responses. (left). Cb3a bipolar cell responses from all stimulation epochs were averaged into three groups. The black trace shows the average bipolar cell response in the absence of a cone tpsc. The red trace shows the average bipolar cell response when the cone released one or more transmitter quanta and the bipolar cell was simultaneously categorized as failing to respond. The green trace shows the average bipolar cell response when the cone released 1 or 2 quanta while the bipolar cell was categorized as responding. The green trace illustrates the expected time course of the epsc. Number of traces averaged in parentheses. Same pair as Fig 3A,B. (right). Similar analysis for a cb3b cell. The red trace averages suggest that categorized response failures in cb3 cells do not consistently contain small epscs. The original amplitude criterion for judging a failure versus a success was based on trace noise (see methods). We cannot exclude that further steps to reduce noise or optimize filtering (*e.g.*, sylgarding the pipette, blocking voltage-dependent membrane currents, matched filter detection) would reveal that some traces judged to be failures might contain small events.

**Figure S4:** Analysis of GluK1 and cb3 cell coverage at cone synapses. **A.** (*i*) Larger image from which Fig 3G, left was obtained. GluK1 is shown in green (Scale = 2 µm). (*ii*) Binarized image with squares centered around selected terminals. (*iii*) Examples of individual, binarized GluK1 labeling. (*iv*) Average of individual squares. (*v*) 2D power spectrum for determining filter cut-off. Components with power less than 55.56 dB were removed. (*vi*). Fourier-filtered image of the average from *iv* showing contour lines. Gray scale is from 0 to 255. (*vii*) Ellipses were fitted to outer and inner profiles using a Hough transform. (*viii*) Location of the unweighted centroid (green circle). (*ix*) Location of the weighted centroid (red) and the displacement vector (green arrow). Scale bar for the inset is 20 nm. **B.** ($1^{st}$ row) Binarized GluK1 labeling from 4 additional images. ($2^{nd}$ row) Average profiles. ($3^{rd}$ row) Filtered profiles with the filter cut-offs appended. ($4^{th}$ row) Elliptical profiles and centroid displacements. Percentages refer to the average gray scale intensity within the ring. **C.** Selected cb3 cell dendritic terminals excised, averaged, and analyzed as in B.

**Figure S5: A.-C.** Scatterplots for 3 additional cone to cb1a cell pairs. Power law fits (black curves where feasible in the cone-linear range), exponents, and corresponding data points in Fig 4E are indicated. Insets show the maximal cone tpsc and bipolar cell epsc. **D.** Analysis of cb1a cell response failures for evidence of small epscs. (top). Cb1a from Fig 4E (black circles). (bottom). cb1a from Fig 4E (red circles). A deviation from baseline following the stimulus suggests (arrow) that some responses classified as failures may contain small epscs. **E**. A second example of labeled cb1a bipolar cell analyzed for the central location of its contacts at the cone terminal. Confocal image of the bipolar cell in cross section (green labeling at top shows Ribeye). **F.** STED image in the whole mount orientation of the GFP labeled bipolar cell (red), Ribeye (magenta), and GluK1 (green). Terminal profiles are outlined in green. **G.** Placement of terminal (yellow) and contact (cyan) centroids. **H.** Contact centroids were displaced from the terminal centers by a normalized micron distance of 0.175 ± 0.126 (different from 0, p = 0.0031). The central contacts occupied 3.1 ± 2.1% of the terminal area.

**Figure S6:** Correlated responses in postsynaptic bipolar cells during depolarization of a common presynaptic cone. **A**. Trial to trial variations in the simultaneous responses of a cb1a and cb2 cell (data from the pair in Fig 5B) during a single epoch of cone depolarization. The epsc peak values of the cb1a cell were scaled by y = 190 + 5.7x and superimposed on the values for the cb2 cell. *(inset)* The responses had a correlation coefficient, $r^2$, of 0.63 (point color corresponds to Fig 5B; $r^2$ is scale independent). **B.** Data from the cb1a and cb2 cell pair in Fig 5C. The peak epsc responses in the cb1a cell were scaled by y = 90 + 0.8x. **C.** Data from the cb3b and cb2 cell pair in Fig 5D. The cb3b cell peak responses were scaled by y = 40 + 1.2x. **D.** Data from the cb3a and cb2 cell pair in Fig 5E. The cb3a cell peak responses were scaled by y = 31 + 2.6x. Assuming that the release at neighboring invaginations is uncorrelated, simultaneous responses in the postsynaptic bipolar cells should be uncorrelated if sampling is exclusively from different ribbons but completely correlated if sampling is from the same set of ribbons. The range of $r^2$ values from 0.63 to 0.79 is consistent with overlap in sampling of release sites in a cone terminal.

**Figure S7:** Responses of cb1a and cb3 cell somas during rapid perfusion with low concentrations of glutamate (12.5 – 100 µM) with 18 mM as a reference. **A.** (left) Confocal image of a cb3a cell remnant (magenta) with an axon ramification below the upper ChAT band (green). (right) Responses to concentrations of rapidly applied glutamate. *Inset.* Initial responses shown on expanded amplitude and time bases. Black trace, above, shows the open-tip timing of the solution exchange and include a slight "bounce" due to perfusion pipette oscillation a few milliseconds after the end of the command step. **B.** Responses of cb1a cell soma (cell remnant, magenta, Ribeye label, green) to glutamate concentration steps ranging from 20 to 80 µM with an 18 mM step as a reference. **C.** Peak response versus concentration plot for the data in A. Fit is to a Hill equation with an $EC_{50}$ of ~377 µM and a coefficient of 0.98. For the fit, the baseline was fixed at 0 and the maximum was fixed at 260 pA. Although missing data points in the mid-range, the fit parameters were consistent with those shown in Fig 6D. **D.** Log-log plot of peak response versus glutamate concentration for 4 somas. Circles in A,B indicate correspondence. The linear fit slope gives a Hill coefficient of ~1. The relative glutamate sensitivities of cb3 and cb1a receptors are maintained at low concentrations of glutamate equivalent to those occurring at basal contacts during quantal release. The $IC_{50}$ was measured in a cb3a (**E**) and a cb1a cells= (**F**) by exposing a soma to a continuous steam of glutamate at concentrations between 0 and 40 µM. Steady-state inhibition was measured during a rapid shift into a solution that contained 18 mM glutamate. The time course of the solution change in E was measured with an open pipette tip. The time course was estimated in F. Cell remnants were counterstained with an antibody to ChAT (left). **G.** Plots of peak response versus steady glutamate concentration for the cells in E and F were fitted with a Hill curve to obtain the $IC_{50}$. An anomalously low peak response in 0.32 µM glutamate was omitted from the cb3a cell fit. **H.** Aggregate $IC_{50}$ measurements (cb1a: $IC_{50}$ = 30.4 ± 10.2 µM, Hill Coef. = 1.9 ± 1.0, n = 7 somas, mean ± S.D.; cb3: $IC_{50}$ = 8.3 ± 8.2 µM, Hill Coef. = 2.8 ± 2.2, n = 8 somas, 5 cb3a and 3 cb3b; means are different p = 0.0004). **I.** Paired pulse responses to 60 ms applications of 18 mM glutamate. **J**. Paired-pulse recovery plots were normalized to the maximum response and averaged. Recovery was fitted with a single (cb3 somas) or double (cb1 somas) exponential curve. All experiments were performed in the presence of 35 µM GYKI53655.

**Figure S8**: Cb1a cell receptor and synapse modeling. **A.** Rate constants in a 9-state receptor model (Hausser and Roth, 1997) were obtained by Particle Swarm Optimization using the results from Fig 6 and Fig S7 (Supplemental Table 3). **B.,C.** Model and actual responses were compared with respect to the response profile during steps into different concentrations of glutamate, $EC_{50}$ and $IC_{50}$ plots, and the rate of recovery from desensitization. Results from the model had an $EC_{50}$ = 1.78 mM, an $IC_{50}$ = 41.7 μM, and recovery $\tau_{fast}$ = 68.9 ms (49%) and $\tau_{slow}$ = 884.6 ms. **D.** Diagram of the synapse model constructed in Cellblender from different perspectives. The presynaptic surface is impermeable. The postsynaptic has a zone (blue) where glutamate can leak out of the 16 nm cleft. 10% of the zone's surface area is space where the rectangular solids represent the dendrites of basally contacting bipolar cells. **E.** Plot of glutamate concentration as a function of time at various distances from a release site obtained from a Monte Carlo simulation. Receptors and transporters were absent for this simulation.

**Figure S9.** Analysis of vesicle density on ribbons. **A.** Electron micrograph showing a vertically oriented cone with the terminal outlined (cyan). The terminal base has a concave profile. **B.** Tangential section through the cone terminal region showing two profiles (cyan). The upper terminal is sectioned through the apex concavity and shows a cluster of ribbons. **C.** Lower terminal in **B** at higher magnification. The section captures the cone near the base of the pedicle. The central region is dominated by postsynaptic dendrites. The dashed lines highlight the major dendritic branches. **D.** An image used to estimate ribbon length. **E., F.** Sample micrographs used to determine ribbon height and vesicle density on the ribbon (D-F, are presented at the same scale). Arrows in E denote a ribbon that was section vertically relative to the arciform density and synaptic triad. Arrows and dashed line in F denote a ribbon with vesicles aligned along one side. **G.** Shows two ribbons, captured in a vertical section, that were used to estimate ribbon length. **H.-J.** Histograms of ribbon length, ribbon height, and the distances between SVs on the ribbon. **K.** Histogram of vesicle diameters.

**Figure S10:** cb1a cell responses demonstrate a non-linearity when the amount of transmitter release was reduced by increasing cone stimulus frequency. Increases in frequency reduce the RRP and thus transmitter released. Cones were stimulated in the loose seal mode. Stimulus (1 ms) strength was adjusted to produce a maximal epsc response during a 1 Hz train. **A.** Select trains showing epsc responses in cb2, cb1a, and cb3b cells with the indicated repetition rate. **B.** Plots of peak response versus trial number. Trains that did not produce a response after the initial pulses were omitted for clarity. **C.** Plots of the normalized response (average of the final 8 responses in the train divided by the first response) as a function of interpulse interval for the cb2, cb1a, and cb3b cells. The rightmost plot shows aggregate data. The cb2 results in the first column are replotted from Grabner et al, 2016.

#### Supplementary Movies

**Movie S1:** 3D STED microscopic reconstruction of a retinal whole mount labeled for PSD95 (red) and EAAT5 (cyan).

**Movie S2:** 3D STED microscopic reconstruction of a retinal whole mount labeled for GFP in a cb2 cell (red), GluA4 (yellow), and Ribeye (magenta).

**Movie S3:** 3D STED microscopic reconstruction of a retinal whole mount labeled for GFP in a cb3b cell (red), GluK1 (green), and the overlap region (white).

**Movie S4:** 3D STED microscopic reconstruction of a retinal whole mount labeled for GFP in a cb1a cell (red), GluK1 (green), Ribeye (magenta), and GluA4 (white).

Figure S1

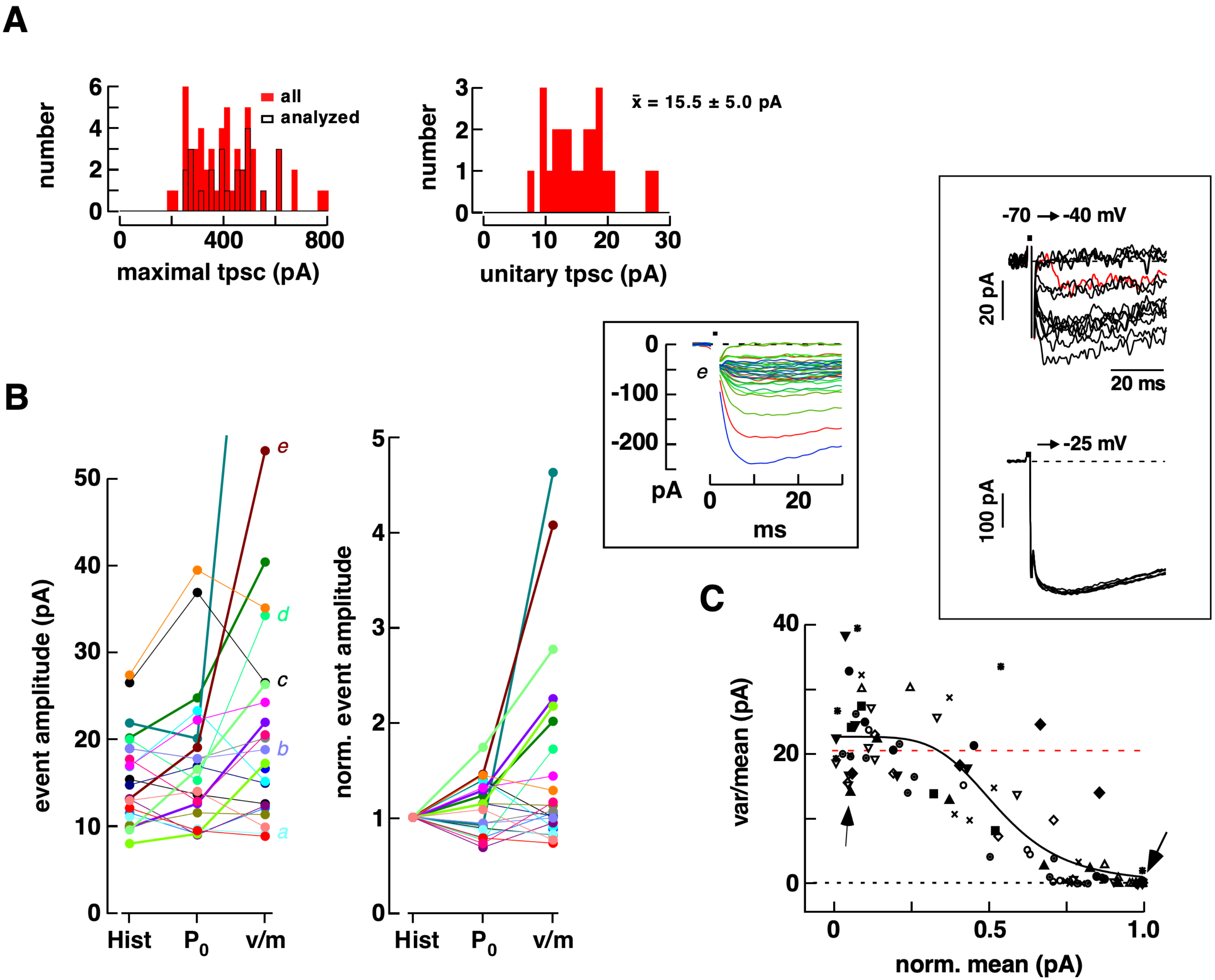
A
number
all
analyzed
maximal tpsc (pA)
unitary tpsc (pA)
x̄ = 15.5 ± 5.0 pA
-70 → -40 mV
20 pA
20 ms
→ -25 mV
100 pA
B
event amplitude (pA)
e
d
c
b
a
Hist
P0
v/m
norm. event amplitude
pA
ms
C
var/mean (pA)
norm. mean (pA)

Figure S2

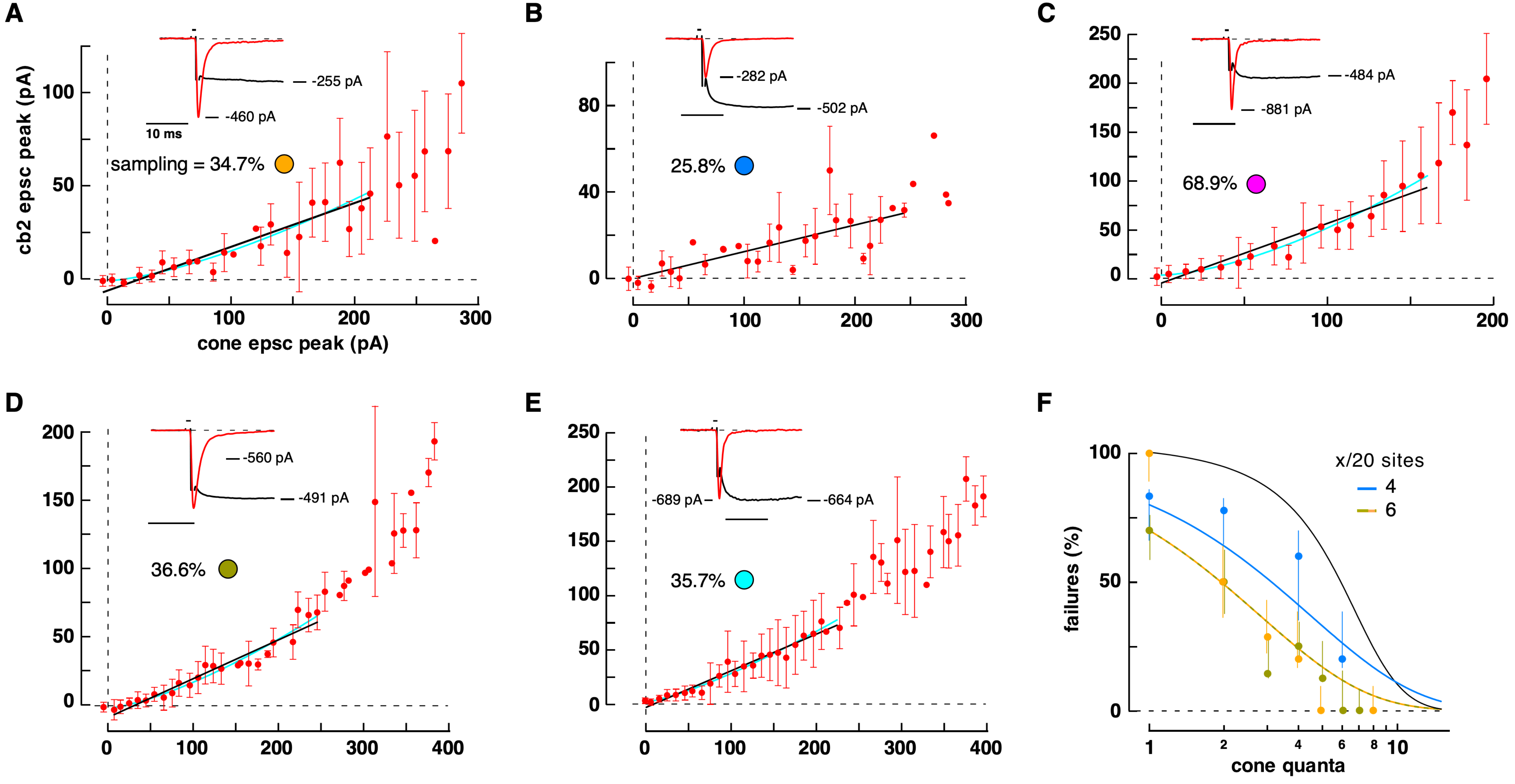

A
cb2 epsc peak (pA)
cone epsc peak (pA)
-255 pA
-460 pA
10 ms
sampling = 34.7%
B
-282 pA
-502 pA
25.8%
C
-484 pA
-881 pA
68.9%
D
-560 pA
-491 pA
36.6%
E
-689 pA
-664 pA
35.7%
F
failures (%)
cone quanta
x/20 sites
4
6

### Figure S3

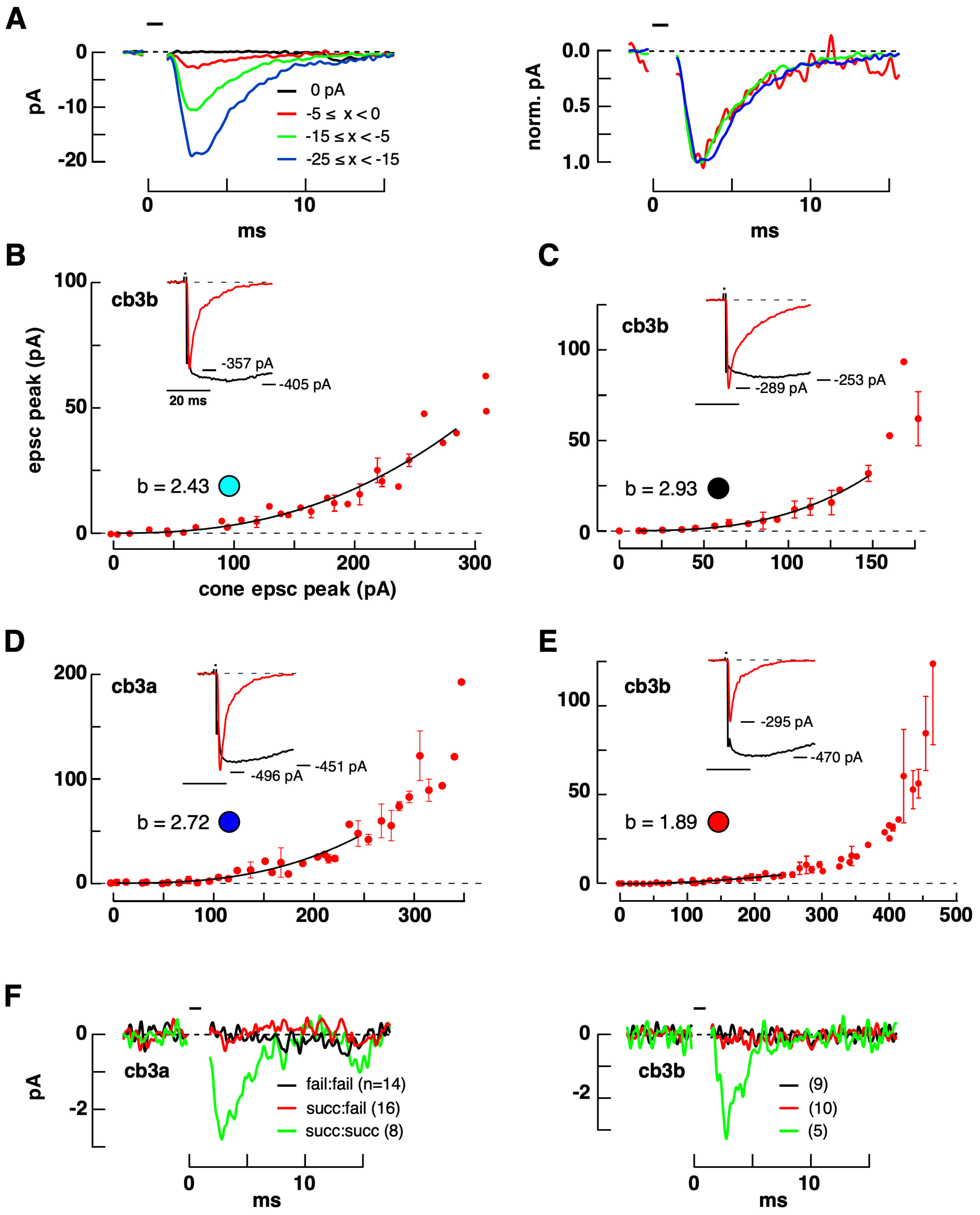
A
pA
0
-10
-20
0 pA
-5 ≤ x < 0
-15 ≤ x < -5
-25 ≤ x < -15
0
10
ms
norm. pA
0.0
0.5
1.0
0
10
ms
B
cb3b
epsc peak (pA)
100
50
0
-357 pA
-405 pA
20 ms
b = 2.43
0
100
200
300
cone epsc peak (pA)
C
cb3b
100
50
0
-289 pA
-253 pA
b = 2.93
0
50
100
150
D
cb3a
200
100
0
-496 pA
-451 pA
b = 2.72
0
100
200
300
E
cb3b
100
50
0
-295 pA
-470 pA
b = 1.89
0
100
200
300
400
500
F
cb3a
pA
0
-2
fail:fail (n=14)
succ:fail (16)
succ:succ (8)
0
10
ms
cb3b
0
-2
(9)
(10)
(5)
0
10
ms

### Figure S4

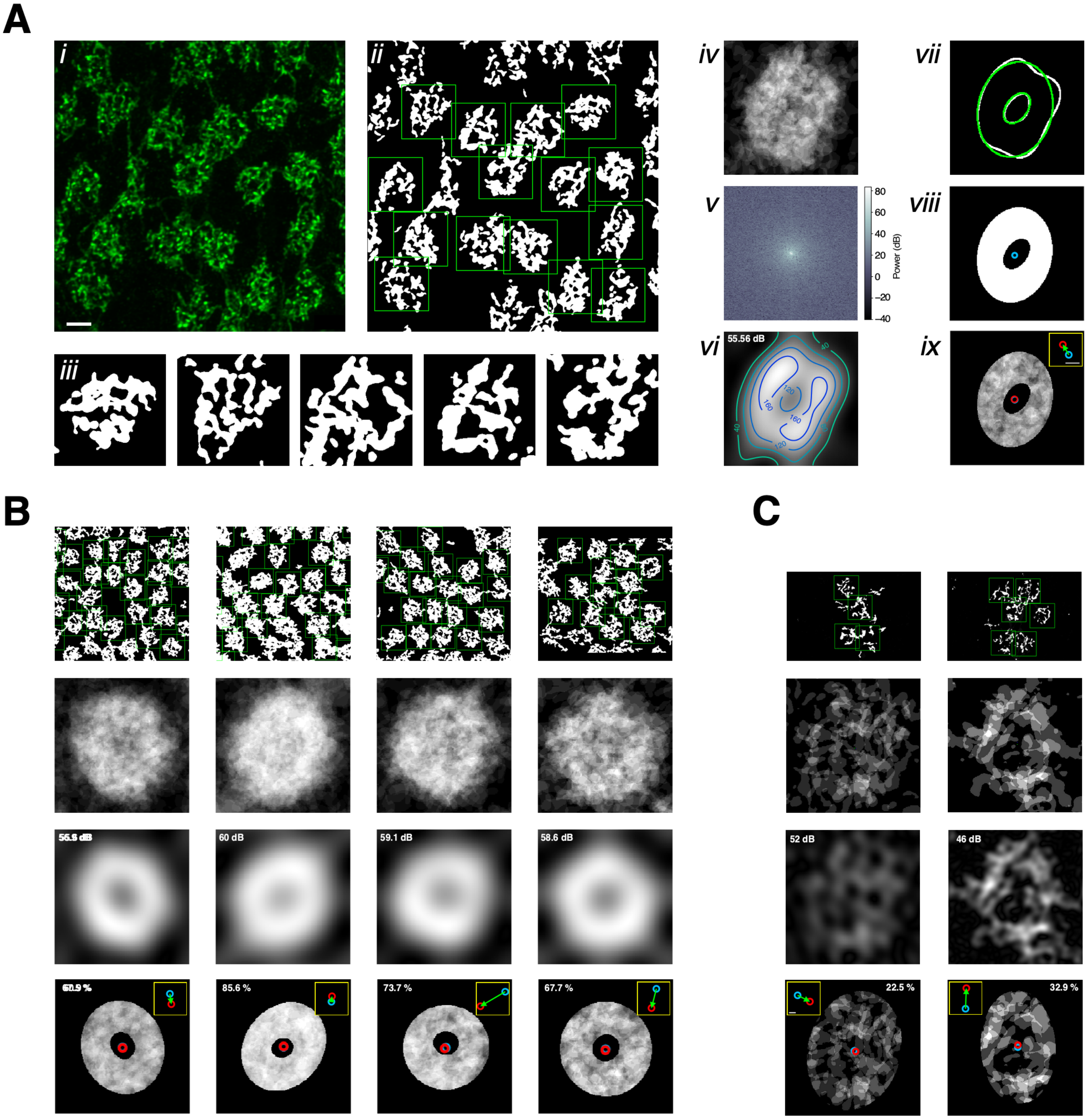
A
i
ii
iii
iv
v
vi
vii
viii
ix
Power (dB)
80
60
40
20
0
−20
−40
55.56 dB
B
C
60 dB
59.1 dB
58.6 dB
85.6 %
73.7 %
67.7 %
52 dB
46 dB
22.5 %
32.9 %

Figure S5

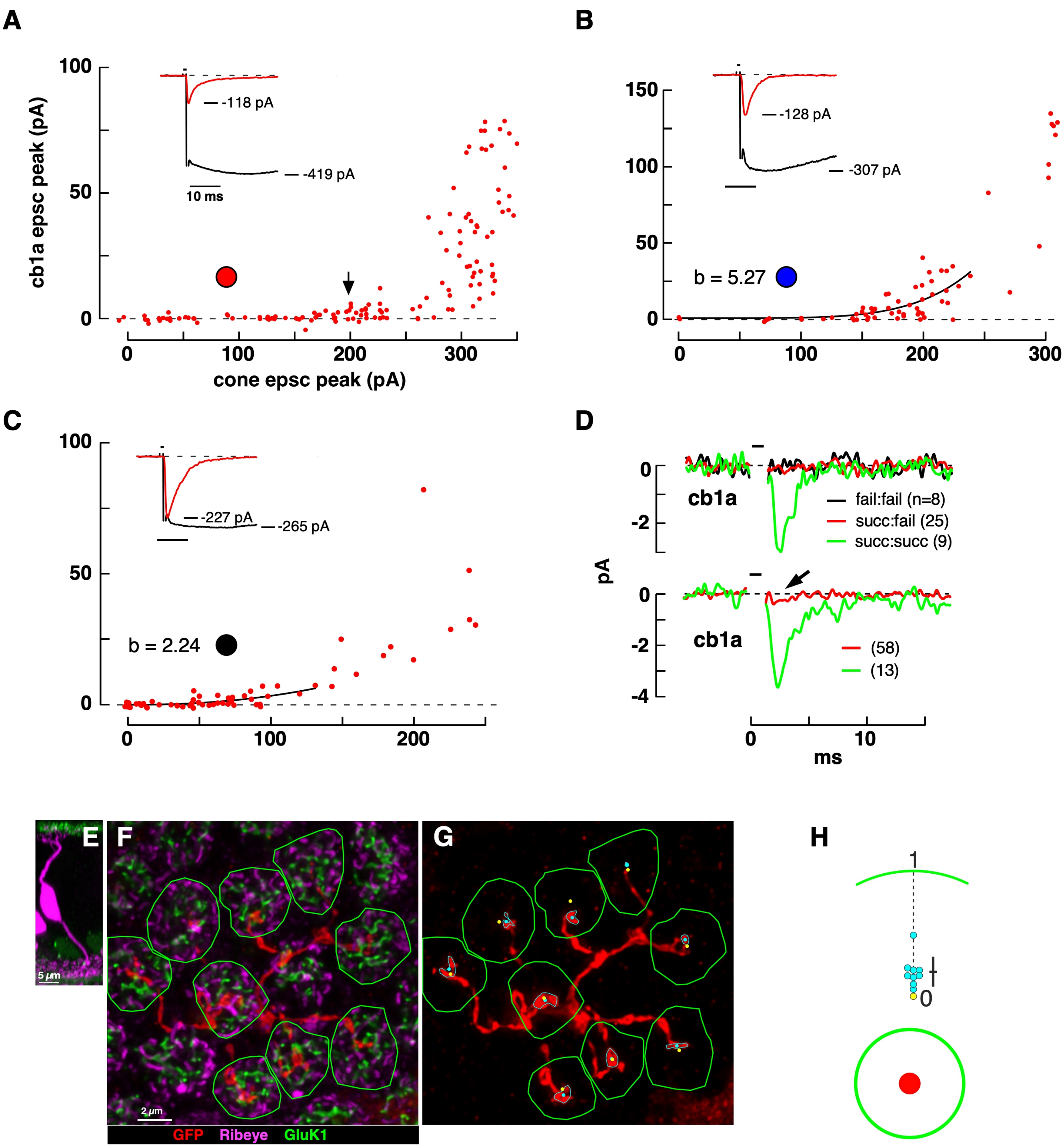

A
100
50
0
cb1a epsc peak (pA)
-118 pA
-419 pA
10 ms
0
100
200
300
cone epsc peak (pA)
B
150
100
50
0
-128 pA
-307 pA
b = 5.27
0
100
200
300
C
100
50
0
-227 pA
-265 pA
b = 2.24
0
100
200
D
0
-2
cb1a
fail:fail (n=8)
succ:fail (25)
succ:succ (9)
pA
0
-2
-4
cb1a
(58)
(13)
0
10
ms
E
5 µm
F
2 µm
GFP Ribeye GluK1
G
H
1
0

Figure S6

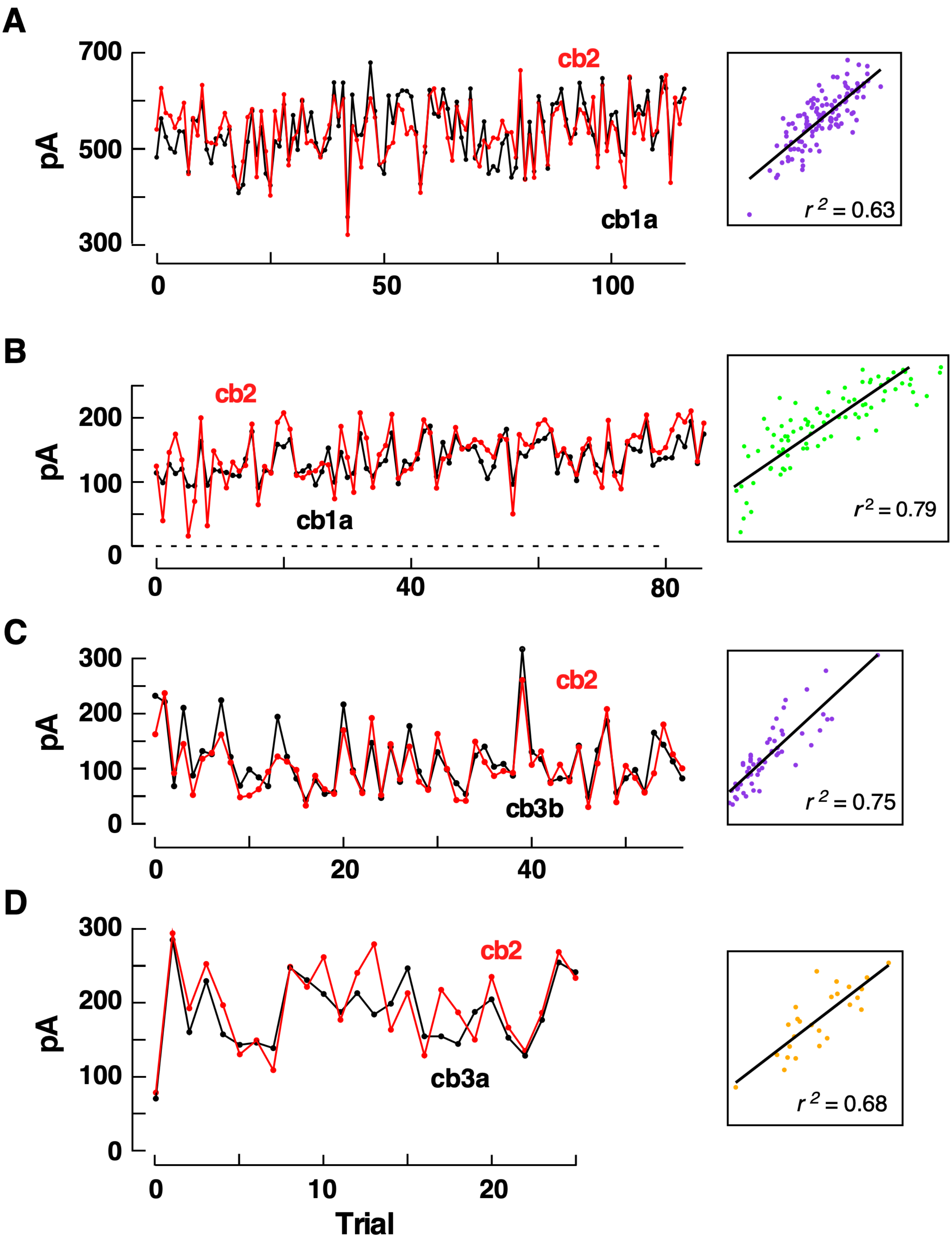
A
700
500
300
pA
cb2
cb1a
0
50
100
r 2 = 0.63
B
200
100
0
pA
cb2
cb1a
0
40
80
r2 = 0.79
C
300
200
100
0
pA
cb2
cb3b
0
20
40
r 2 = 0.75
D
300
200
100
0
pA
cb2
cb3a
0
10
20
Trial
r 2 = 0.68

Figure S7

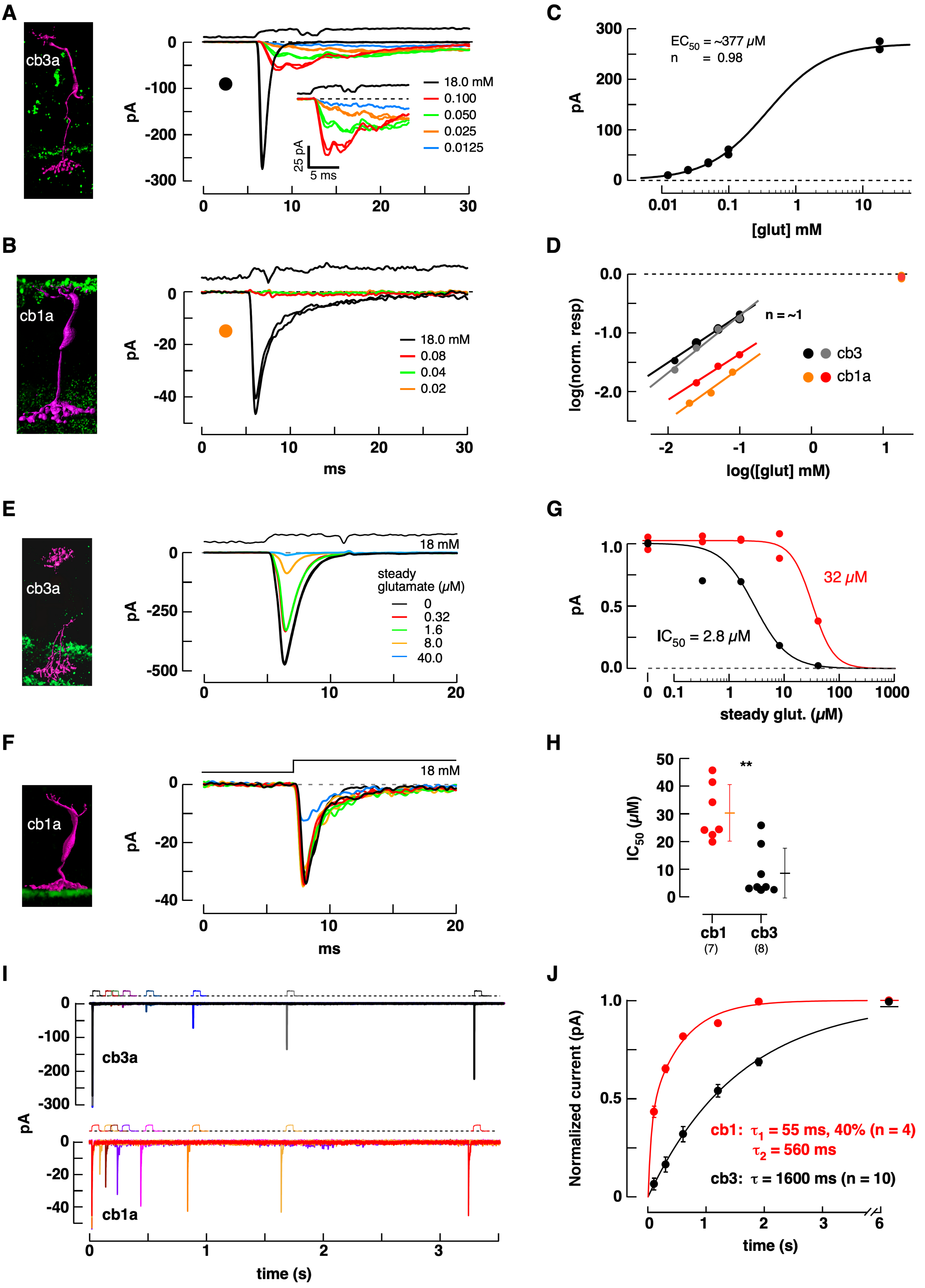

A
cb3a
pA
0
-100
-200
-300
0
10
20
30
18.0 mM
0.100
0.050
0.025
0.0125
25 pA
5 ms
C
300
200
100
0
pA
EC50 = ~377 μM
n = 0.98
0.01
0.1
1
10
[glut] mM
B
cb1a
pA
0
-20
-40
18.0 mM
0.08
0.04
0.02
ms
D
0.0
-1.0
-2.0
log(norm. resp)
n = ~1
cb3
cb1a
-2
-1
0
1
log([glut] mM)
E
cb3a
18 mM
steady
glutamate (μM)
0
0.32
1.6
8.0
40.0
pA
0
-250
-500
20
G
1.0
0.5
0.0
32 μM
IC50 = 2.8 μM
100
1000
steady glut. (μM)
F
cb1a
18 mM
H
50
40
30
IC50 (μM)
**
cb1
(7)
cb3
(8)
I
cb3a
cb1a
time (s)
J
Normalized current (pA)
cb1: τ1 = 55 ms, 40% (n = 4)
τ2 = 560 ms
cb3: τ = 1600 ms (n = 10)
6
time (s)

Figure S8

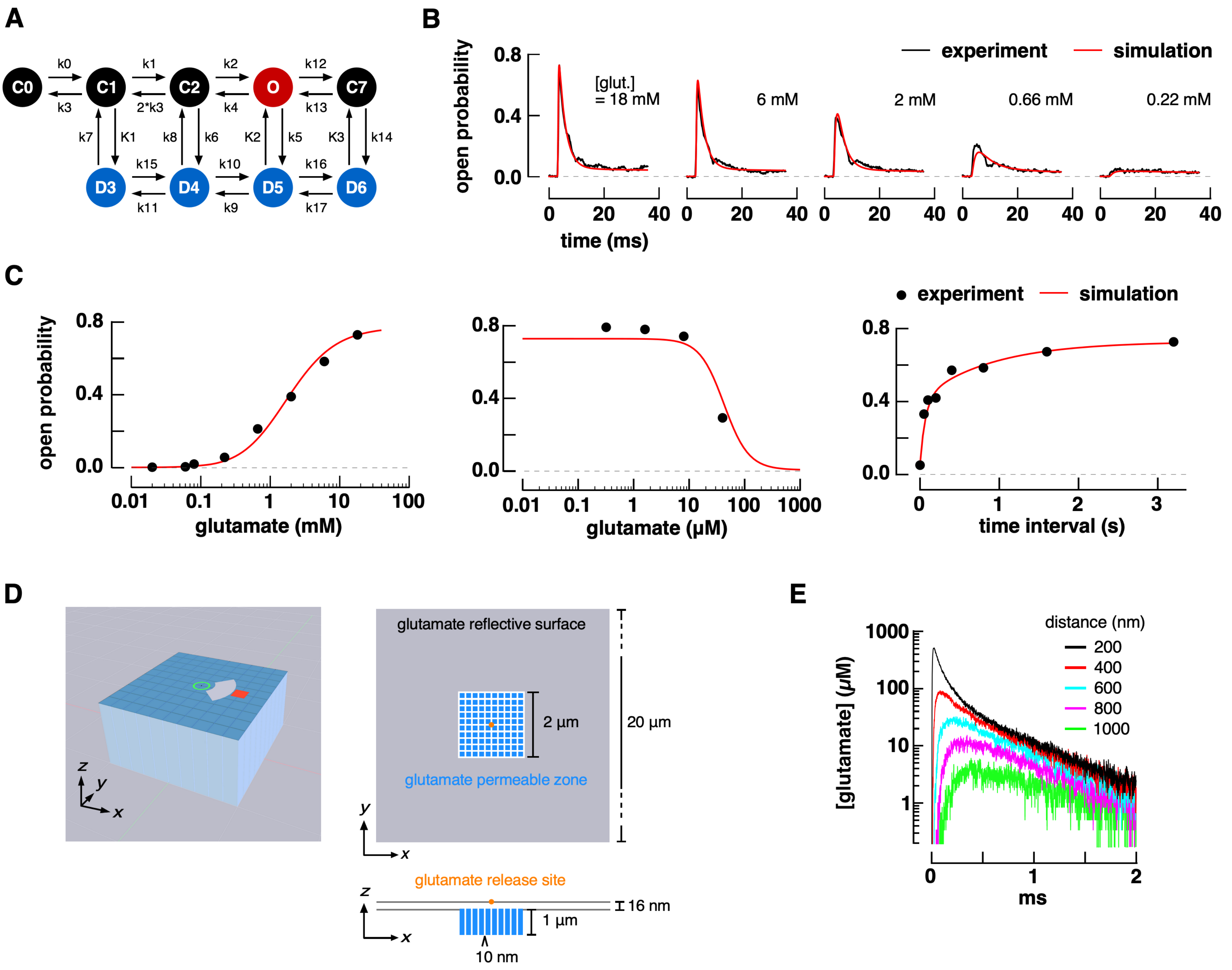

A
C0
C1
C2
O
C7
D3
D4
D5
D6
k0
k3
k1
2*k3
k2
k4
k12
k13
k7
K1
k8
k6
K2
k5
K3
k14
k15
k11
k10
k9
k16
k17
B
experiment
simulation
open probability
[glut.]
= 18 mM
6 mM
2 mM
0.66 mM
0.22 mM
time (ms)
C
glutamate (mM)
glutamate (μM)
time interval (s)
D
glutamate reflective surface
2 μm
20 μm
glutamate permeable zone
glutamate release site
16 nm
1 μm
10 nm
E
distance (nm)
200
400
600
800
1000
[glutamate] (μM)
ms

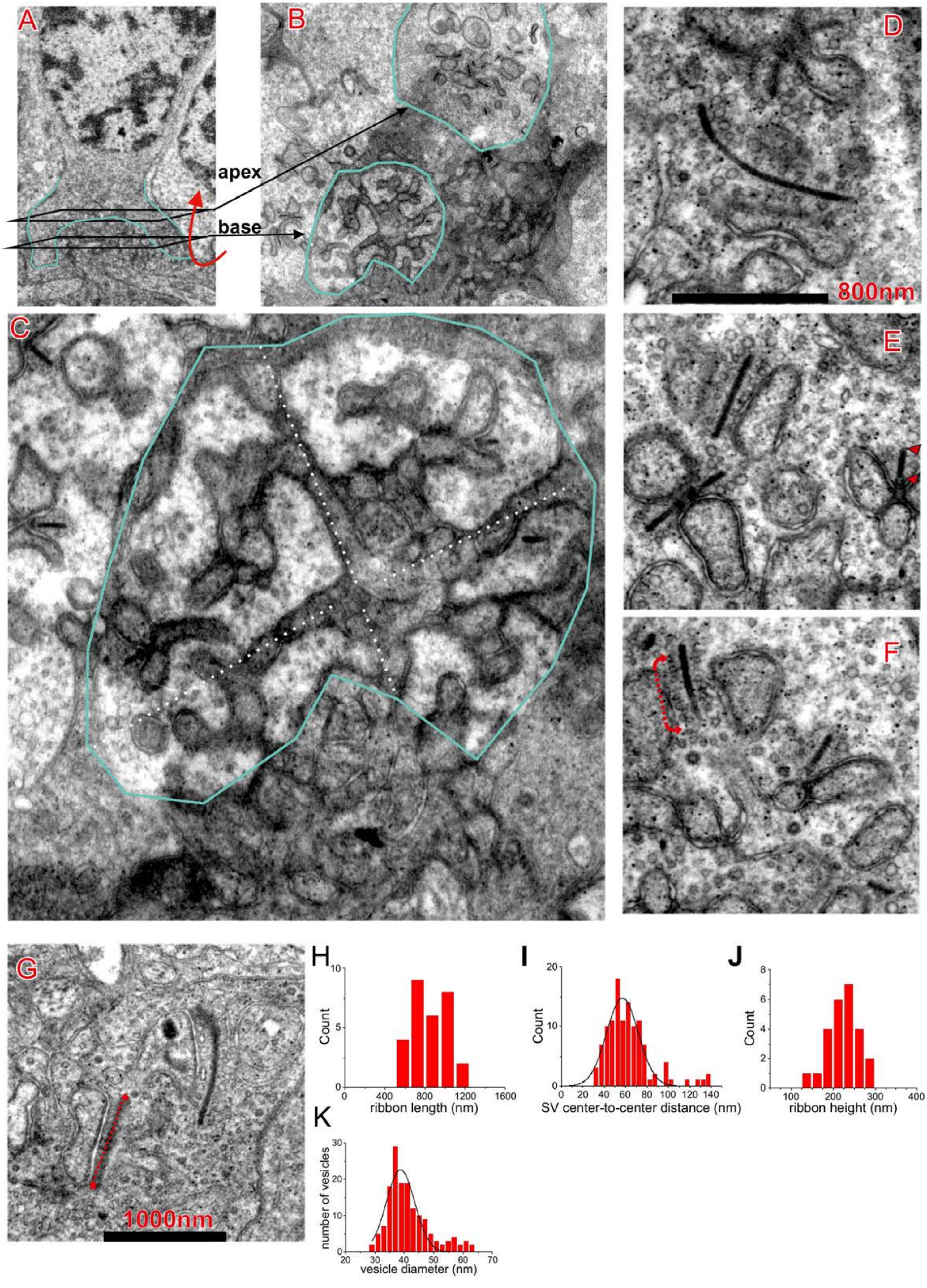

A
B
D
apex
base
800nm
C
E
F
G
1000nm
H
Count
ribbon length (nm)
I
Count
SV center-to-center distance (nm)
J
Count
ribbon height (nm)
K
number of vesicles
vesicle diameter (nm)

Figure S10

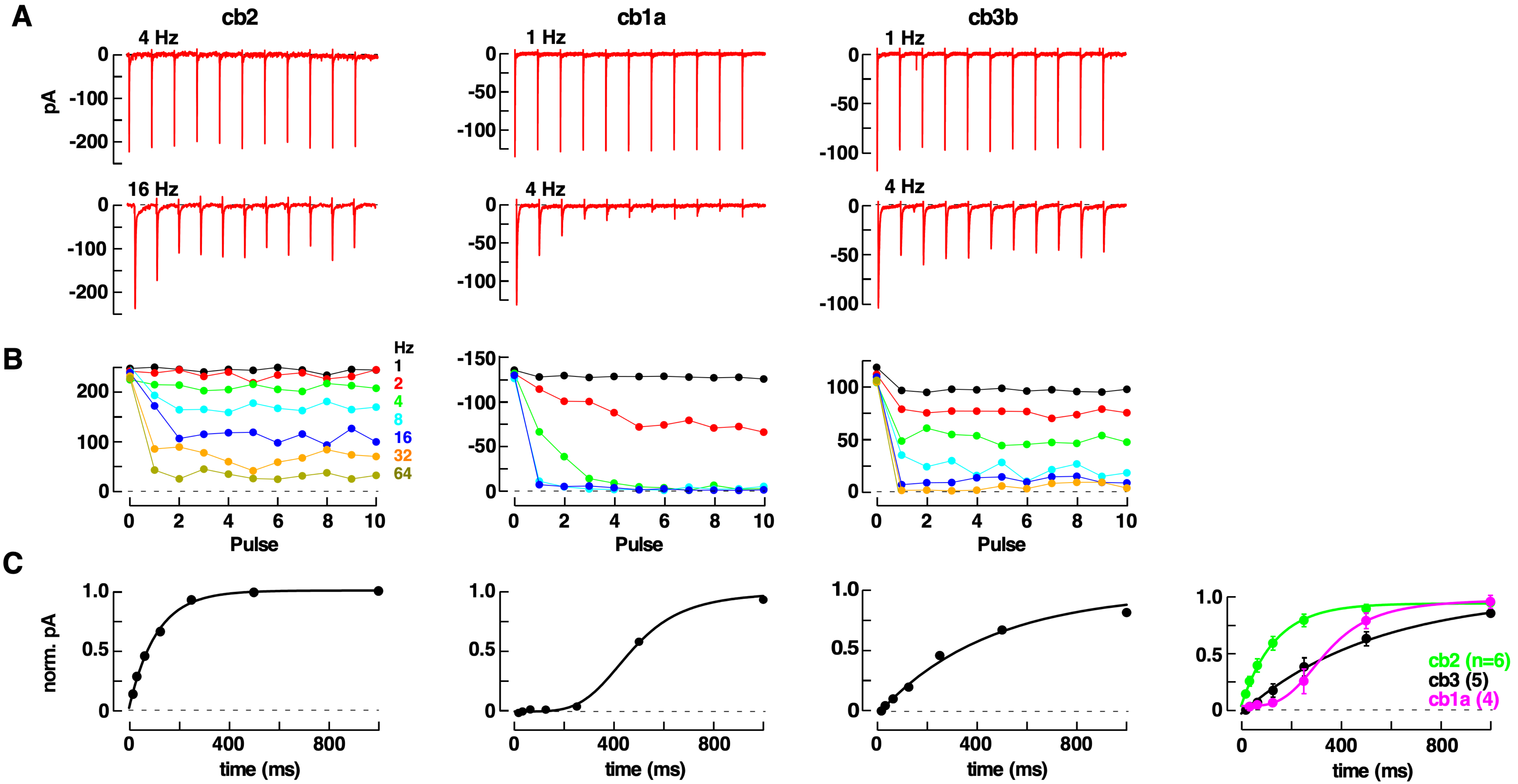

A
cb2
cb1a
cb3b
4 Hz
16 Hz
1 Hz
4 Hz
1 Hz
4 Hz
pA
B
Hz
1
2
4
8
16
32
64
Pulse
Pulse
Pulse
C
norm. pA
time (ms)
time (ms)
time (ms)
time (ms)
cb2 (n=6)
cb3 (5)
cb1a (4)
